# Self-organisation and persistence of antibiotic resistance in evolving plasmid communities

**DOI:** 10.1101/2020.09.15.297721

**Authors:** Martin Zwanzig, Uta Berger

## Abstract

A key source of genetic variation of microbial populations are plasmids: extrachromosomal genetic elements that replicate autonomously and can be highly mobile between individual cells. Diverse plasmids were found in environmental samples and bacterial populations. Here we explore the mechanisms that help to preserve this gene pool as a fundamental basis for bacterial adaptation. An individual-based model of the plasmidome is presented and used to investigate how intra- and intercellular competition between diverse plasmid types affects the evolution of plasmid communities. It indicates the relative importance of stochastic and deterministic drivers of plasmid persistence both under neutral conditions and when the environment selects for specific plasmid-encoded traits such as antibiotic resistance for a certain period of time. We found that evolving plasmid communities exhibit a cyclical dynamics that contributes to the maintenance of plasmid diversity and the persistence of costly plasmid-mediated antibiotic resistance after stopped abiotic selection.

## Introduction

Bacteria are often exposed to environmental changes that require rapid genetic adaptation. Access to the relevant genes can therefore be crucial for bacterial survival. In addition to the stable core genome, which can represent for example around 2000 genes for a single bacterial cell (assessed for 20 *Escherichia coli* strains)^1^, bacteria gain and lose genes, which can be highly mobile between bacterial cells^2^. These non-core genes increase the diversity of a species’ pan-genome and can make up around 90% of it^3^, where plasmids contribute a large proportion^4^. Although more and more studies provide insights on the plasmidome^4–10^, which refers to the entire plasmid DNA of an environmental sample^9^, the mechanisms that keep plasmid diversity are not fully understood.

Remarkable progress has been made investigating the persistence conditions of plasmids in a single plasmid - single host^11^–17 or single plasmid - multiple host environment^18–22^, but we are just beginning to understand the role of a co-occurrence of multiple plasmid types, either in a single^23–32^ or multiple host^33^ framework. It has been shown that the number, size and functions of the plasmids found in different genera^34^, single species^5,8^ or in environmental samples^4,6,7,9,35–39^ can vary remarkably.

Both ecological and genetic factors were reported to affect plasmid distribution^10^. Many plasmids encode selfish traits as stability and conjugation^40^, which improve their inheritance in a cell line or allow the infection of new bacterial hosts. Considering pairwise plasmid-host interactions, high conjugation rates^11^ as well as high or low frequency pulses of positive selection of accessory functions such as antibiotic resistances^15^ can slow down or prevent plasmid loss. Positive selection can also help to maintain non-transmissible plasmids, since a high frequency promotes compensatory evolution to ameliorate the plasmid cost^14^. Another study^41^ found that a small non-mobile plasmid encoding for antibiotic resistance can be stable in its host over evolutionary timescales in the absence of antibiotics. The results of this study indicate that the resolution of transcription-replication conflicts are likely to improve the efficiency of plasmid inheritance in a way that allows plasmids to reach fixation similar to neutral alleles. Considering multiple hosts, conjugative plasmids may only be stably maintained in one of the hosts, but this can act as a reservoir for other hosts that become continuously infected^42^. Such source-sink dynamics might be affected considering multiple plasmid types, since intracellular interactions between two^31^ or three^32^ distinct conjugative plasmids can affect the conjugation efficiency, with a trend towards a reduction of the conjugation rate.

Bioinformatic analysis revealed that co-infection with multiple plasmids is common across a wide range of bacterial phyla^29^. For example, the large horizontal gene pool of rhizobia indicates that the pool of plasmids rather than an individual plasmid is stable^43^. Such plasmidomes can contain a huge number of diverse plasmids that span multiple plasmid incompatibility groups^44^. Incompatibility is referred to the inability of plasmids to stably co-exist in the same host cell line due to similarities in the replication and partition systems^34^. There are, for example, 27 Inc groups in *Enterobacteriaceae*. Meta-analysis of sequence data further reported that associations between small and large plasmids are specifically frequent^29^, which has also been found in ecological communities^10^, but it is still not completely clear why. It also remains unsolved, how a high diversity of plasmids that encode no or unknown accessory functions persist, for example, in freshwater habitats^7^.

Understanding how species interactions modulate biodiversity is a central aim in ecology that benefits from a recognition of the multidimensional nature of a species niche^45^. It has been found that interactions between trophic levels can determine the survival of a resistance plasmid more than lethal antibiotic concentrations^46^, whereas another study reported significant geospatial and temporal variation of resistance loads, with a strong correlation to the number, proximity, size and type of the surrounding wastewater treatment plants^47^. Although the relative importance of single factors and the details of mechanisms controlling community structure, succession and biogeography are still a central debate, patterns of biodiversity are likely simultaneously governed by deterministic and stochastic processes^48,49^. This reflects a new theoretical framework combining traditional niche theory with neutral theory, which relates community structure to deterministic factors such as environmental conditions and the traits and interactions of species as well as to stochastic processes such as birth, death, colonization, extinction and speciation.

Plasmid dynamics is a multi-scale phenomenon, because individual plasmids interact with each other on the same scale when they are in the same host cell, but they also interact with their host cell and with other host cells and their plasmids. Such complex systems are inherently limited in their predictability, because interactions between system components can generate new information that determine the systems future^50^. One of the fundamental questions is how interactions of the systems components regulate its internal states, e.g. through feedback-loops, which can trigger the self-organization of initially disordered complex systems.

The work presented here is based on a prior study^30^ that demonstrated how the interaction between one type of conjugative and one type of non-transmissible plasmid can promote plasmid persistence. Here, we extended this approach and compiled information relevant for a description of the interaction of diverse plasmid types in a clonal, surface-attached bacterial population by means of an individual-based simulation model of the plasmidome. It aims to investigate the role of interactions between diverse plasmids on plasmid community dynamics and the fate of antibiotic resistance. In brief, the model considers that plasmid types spread spatially by cell division and horizontal transfer (conjugation) with respect to their individual plasmid costs associated with replication, conjugation and accessory genes (see Figure 1a for the general concept, Figure 1-figure supplement 1 for the characteristics of the initial plasmid community, Figure 1-figure supplement 2 for a flow chart of all processes and Figure 1-figure supplement 3 for the spatial representation of model entities). Plasmids get lost due to competition for space, whereas (i) the host cell’s ability to compete with neighboring bacteria decreases with increasing costs for all resident plasmids, (ii) plasmid types of the same incompatibility group cannot co-reside in the same host cell, and (iii) the host cell can be killed through the action of an antibiotic (if not carrying a resistance plasmid, but antibiotics are present). Robustness tests show that the model is flexible in dealing with a wide range of assumptions. They are presented in the results section after the main findings. A more detailed model description is presented in the methods section.

**Figure 1.**
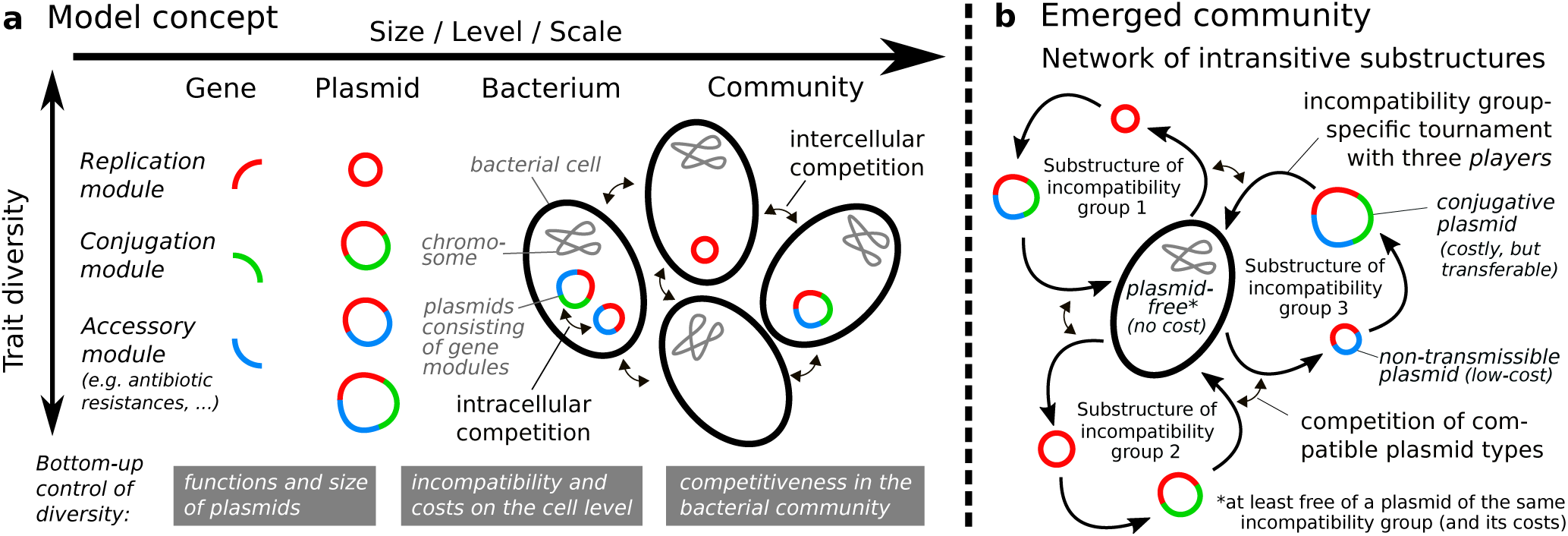
Concept and emerging community characteristics of the individual-based plasmidome model. **a**: Plasmid traits are determined by their gene modules, which are considered to be differently costly for the host, but might also be able to support the plasmid’s propagation, e.g. by conjugation and/or the provision of additional functions such as resistance to antibiotics. **b**: Simulating evolutionary selection reduced plasmid content, but led to the emergence of a stable community, representing a network of intransitive substructures that promote the maintenance of at least one conjugative and one (less costly) non-transmissible (or weakly infectious) plasmid type per plasmid incompatibility group (here illustrated for 3 Inc groups). This is not working in the same way when conjugation is considered to be free of additional costs (see robustness test on “cost of conjugation”).

We performed two different simulation experiments that show the evolution and dynamics of a diverse plasmid community (i) in ‘non-polluted environments’, which refers to the complete absence of abiotic selection, and (ii) in ‘polluted and restored environments’, where the action of an antibiotic that kills sensitive cells stops after some time and no longer benefits costly resistance plasmids.

## Results

To test if at all, how long or to which degree plasmid diversity can be maintained by intra- and inter-cellular interactions alone, we run our model assuming that the environment does not select for any plasmid-encoded trait. The initial plasmid community was constructed by ca. 40,000 different plasmid types, each belonging to one of the given incompatibility groups and associated with random costs for replication (*α*_*rep*_), accessory genes (*α*_*acc*_) and conjugation (*α*_*con*_). Half of the plasmids were considered to be ‘non-transmissible’ (by horizontal gene transfer) and to impose no costs for conjugation.

In a first step, we performed single model runs to assess the basic model behavior when either 1, 3 or 5 different plasmid incompatibility groups are present. Our results show that, although the enormous initial plasmid diversity declines quickly, a small final set of diverse plasmid types is actually maintained in the evolved communities (Figure 2). This is induced by various consequences of the given biotic interactions, which are explained in the following.

**Figure 2.**
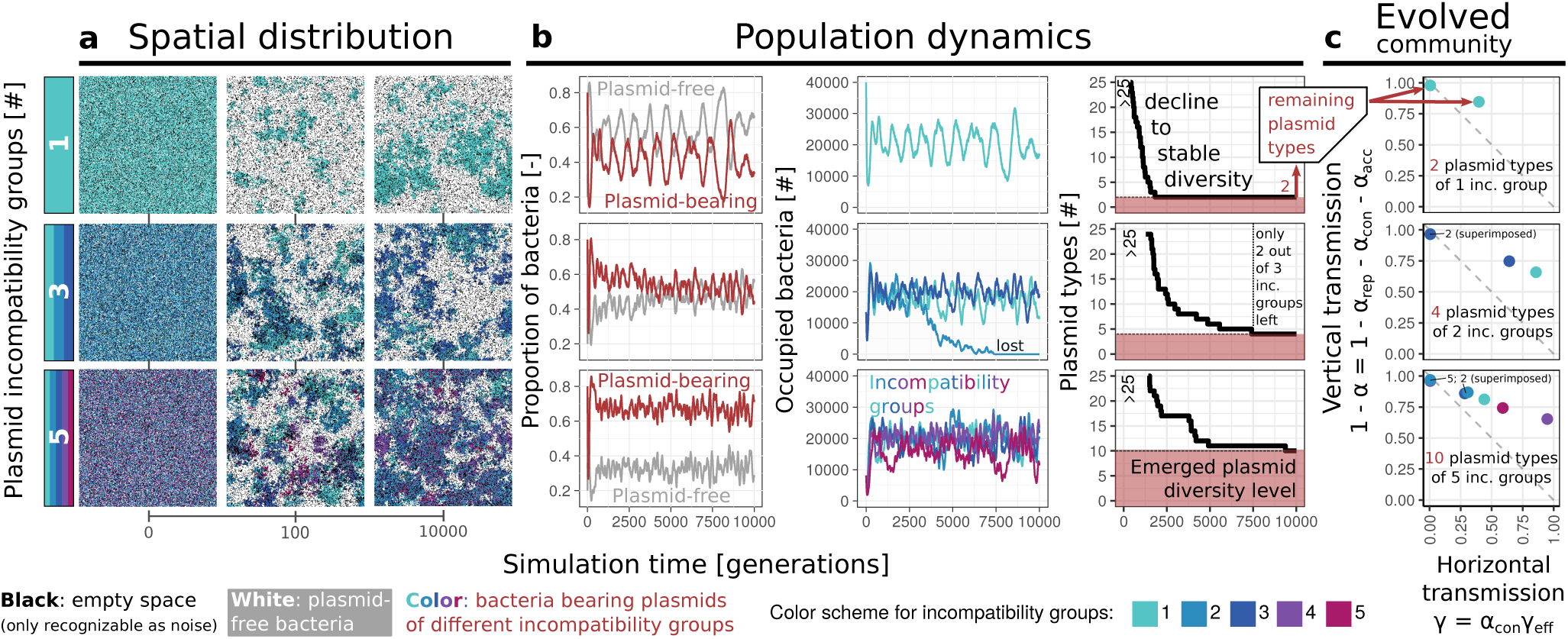
Evolution of plasmid communities. Each row represents a single simulation run considering that the initial plasmid community is composed of either 1, 3 or 5 plasmid incompatibility groups. **a**: The emergence of an aggregated spatial distribution; only incompatibility groups are distinguished, not plasmid types; if multiple incompatibility groups are present in a bacterium, only one color is shown. **b**: Oscillatory population dynamics and the decline of plasmid diversity to a stable state that depends on the number of present incompatibility groups. **c**: Emerged communities comprising one conjugative and one less costly non-transmissible plasmid type per remaining incompatibility group.

### Interactions of neighboring cells influence the spatial distribution of plasmid types

An initially random spatial distribution of diverse plasmid types rapidly evolves to a highly structured form, characterized by an aggregation of plasmid-free bacteria and plasmid types that reside alone or in conjunction with other plasmid types on bacteria (Figure 2a). The more incompatibility groups are present, the greater the number of diverse plasmid types and small-scale changes in their configuration. The proportion of plasmid-free bacteria decreases, the more plasmid incompatibility groups are present in the plasmid community.

### Opposing plasmid lifestyles determine the cyclical dynamics and self-organization of the plasmid community composition

A rapid proliferation of plasmid-bearing cells when plasmid-free cells are common is followed by a fast decline when they are rare (Figure 2b). Oscillations occur due to an enhanced spread of conjugative plasmid types that can access large cluster of recipient cells, which are either plasmid-free or at least free of a plasmid belonging to the same incompatibility group. Surprisingly, however, this horizontal propagation does not necessarily lead to a strong increase in the frequency of these conjugative plasmids (Figure 2-figure supplement 1a). The reason for this is that the host cells of the conjugative plasmid can be simultaneously displaced by other cells with a higher fitness that carry a plasmid of the same incompatibility group. Video 1 shows how cells carrying a less costly non-transmissible plasmid follow the infection front of a conjugative plasmid in an evolved plasmid community - an example for the spatio-temporal pattern that is underlying such dynamics. Thus, the more a conjugative plasmid spreads horizontally, the more cells that carry a plasmid of the same incompatibility group, but have a higher growth rate, can spread as a direct result. In the evolved communities, low-cost non-transmissible plasmids that co-occur with highly infectious conjugative plasmids of the same incompatibility group can therefore show strong fluctuations in their frequency whereas their conjugative counterparts do not. Such an replacement effect is not so strong if the host fitness differences are rather small. When locally available clusters of recipient cells have been infected and the number of successful horizontal gene transfer events drops, the chance for a successful (re-)proliferation of plasmid-free cells or other cells carrying only a reduced number of plasmids is given. As the number of cells that are free of plasmids of a certain incompatibility group increases, the horizontal transfer of conjugative plasmids of this incompatibility group is accelerated again. Previous work^30^ already revealed that a community based of plasmid-free cells, one type of non-transmissible plasmid and one type of conjugative plasmid can exhibit a non-transitive relationship: cells carrying conjugative plasmids spread at the expense of plasmid-free cells but are then outcompeted by cells carrying less costly non-transmissible plasmids, which in turn can be outcompeted by plasmid-free cells. Such a three-species tournament similar to the children’s game rock–paper–scissors has also been shown to allow co-existence in other settings^51^. Although the model presented here considers more than two plasmid types and did not impose any such relation, the emerged community characteristics (Figure 2c) and observed spatio-temporal dynamics are similar. It demonstrates that such non-transitive dynamics can also occur when more diverse plasmid-host associations are present and that this is a typical result of the self-organization of an evolving community of diverse plasmids. If more than one plasmid incompatibility group is considered, the amplitudes of the oscillations can be less pronounced (Figure 2b and Video 2), as such intransitive dynamics can simultaneously occur for each incompatibility group and can be disturbed by the huge number of potential inter- and intra-cellular interactions with other plasmid types (Figure 1b).

### Competitive exclusion minimizes plasmid diversity to a locally sustainable level

Our model is initialized with about 40000 different plasmid types, each of which occurs only once. Some of them are more likely to survive, because they have a very high vertical or horizontal transmission fitness. Besides these deterministic factors, there is a high likelihood of stochastic extinctions. This means plasmid types can be eliminated no matter which vertical and horizontal transmission fitness they have, especially if their abundance is (still) low. The oscillatory dynamics between plasmid-free and plasmid-bearing cells intensifies this eradication process, especially when plasmid diversity is still high and many cells carry multiple or single costly plasmids, which makes them easily replaceable by plasmid-free cells or hosts with less plasmid costs. Even all plasmid types of an incompatibility group can get lost due to competitive exclusion (Figure 2b). This is particularly likely when interactions between plasmids of the same incompatibility group are disturbed by a fragmented spatial distribution of the associated plasmid types. If, for example, plasmids of a certain incompatibility group are lost during cell division, the resulting cell has lower plasmid costs and a higher chance to outcompete all host cells that still carry plasmids of this incompatibility group. This can be a long-term process that might be interrupted when these host cells are (re)infected by a conjugative plasmid belonging to the plasmid incompatibility group that they initially lost through segregation. Most interestingly, even the presence of plasmids from other incompatibility groups can prevent the extinction of (non-transmissible) plasmids that would not be able to survive on their own (see maintenance of non-transmissible plasmid of inc. group 1 by cyclic dynamics when embedded in an evolved community comprising four plasmids from two other incompatibility groups (Video 2) versus constant decrease until extinction in isolation (Video 3). The level at which plasmid diversity finally stabilizes depends on the number of incompatibility groups (see Figure 2b and the robustness test on “Population size and more than five incompatibility groups” below).

[ VIDEO 1 ]

https://youtu.be/bHLsAss0iik

**Video 1. Simulation of the evolution of a plasmid community considering one incompatibility group**. The simulation is based on a default parameterization (Table 2), starting with an initially hyperdiverse and completely mixed plasmid community whose progression is followed over 10,000 generations and which has a mortality or dilution rate of 0.2. Thus, every round there is a probability of 20% that a bacterium and its plasmids are removed from the model domain, leaving (a black colored) empty space that can be recolonized by neighboring bacteria according to their fitness that can be affected by a certain plasmid load (colored plasmids representing (varying) non-transmissible and conjugative plasmids of certain incompatibility groups versus white plasmid-free cells). Please note the monitor that shows the remaining number of diverse plasmids and the ‘tp-pb-relation’-plot depicting their horizontal and vertical transmission fitness similar to Figure 2c.

**Table 1.** Model entities and associated state variables.

| ENTITY | ATTRIBUTE | STATES (/SYMBOL) | MEANING |
| --- | --- | --- | --- |
| **Lattice cell**<br>(/Bacterium) | Location | x-y-coordinates | Lattice position |
|  | Occupation | Empty \| Bacterium | Considered as bacterium or not |
| **Plasmid** | Location | x-y-coordinates | Association with bacterium xy |
| | Replication module costs | $\alpha_{rep}$ | Costs associated with the expression, repair and replication of genes from these single plasmid modules |
| | Accessory module costs | $\alpha_{acc}$ | |
| | Conjugation module costs | $\alpha_{con}$ | |
| | (overall) plasmid burden* | $\alpha = \alpha_{rep} + \alpha_{acc} + \alpha_{con}$ | Overall plasmid costs reducing the host cells growth rate<sup>◊</sup> |
| | Conjugation efficiency | $\gamma_{eff}$ | A factor that is used to relate the costs $\alpha_{con}$ to the real probability $\gamma^{\dagger}$ |
| | Plasmid transfer probability* | $\gamma = \alpha_{con}\gamma_{eff}$ | Probability to perform a transfer attempt<sup>◊</sup> |
| | Plasmid incompatibility group | $\eta$ | Plasmids belonging to the same group cannot co-occur in the same bacterial cell • |

\* Indirect parameters that do only represent an aggregation of other parameter values (as indicated in the table)
◊ Further influenced by the proportion of empty neighboring cells (mimicking the local resource availability)
† when $\gamma_{eff} \geq 1$ , conjugation can theoretically compensate its own costs or, when $\gamma_{eff} \geq \alpha/\alpha_{con}$ , even all plasmid costs
• a simplifying assumption that is subject of a robustness test; in reality they use similar replication or partition systems, which prevents a stable co-occurrence in the same host cell line

**Table 2.** Values assigned to global parameters and state variables during model initialization.

| PARAMETER | VALUE | MEANING |
| --- | --- | --- |
| Mortality $\omega$ | 0.2 | Probability of each bacterium and all its residing plasmids to die (/be removed from the system domain, e.g. according to washout, natural mortality and/or predation) |
| Antibiotic action $\upsilon$ | $0^\diamond$ | Probability that a sensitive bacterium is killed due to the action of an antibiotic |
| Segregation $\tau$ | 0.001 | Probability to inherit a plasmid type only to one of both daughter cells during cell division (in addition to potential effects of incompatibility) |
| Immigration $\varpi$ | $0^\diamond$ | Probability that a lattice cell is replaced by an immigrating bacterium (with random plasmid load) |
| Initial bacterial density $\rho_B$ | $1-\omega$ | Proportion of lattice cells that are occupied by a bacterium at the beginning of a simulation |
| Initial plasmid density $\rho_P$ | 0.8 | Proportion of plasmid-bearing cells of the initial bacterial population (only one plasmid per bacterium is generated) |
| Initial resistance density $\rho_{ARP}^*$ | $0^\diamond$ | Proportion of plasmids conferring resistance to antibiotics |
| Initial $\alpha_{rep}$ distrib. | $0.03 \pm 50\%^*$ | Replication module costs (normal distribution mean ± coefficient of variation $>$ minimum size) of the initial plasmid community |
| Initial $\alpha_{acc}$ distrib. $^*$ | $0 - 0.25$ | Accessory module costs (uniform distribution minimum and maximum) of the initial plasmid community |
| Initial $\alpha_{con}$ distrib. | $0.2 \pm 50\%^*$ | Conjugation module costs (normal distribution mean ± coefficient of variation) of the initial plasmid community |
| Initial $\gamma_{eff}$ distrib. | $2 \pm 50\%$ | Conjugation efficiency (normal distribution mean ± coefficient of variation) of the initial plasmid community |
| Initial # inc. groups | 1, 2, 3, 4, 5 (up to 25) | The number of different plasmid incompatibility groups each plasmid of the initial population can belong to (represent single scenarios) |

\* considering a lower limit: costs $> 0.01$
$^\diamond$ in default model application, otherwise $0 - 1$
\* if antibiotic resistances are considered, those plasmids with nearly mean $\alpha_{acc}$ ( $mean_{\alpha_{acc}} \pm sd_{\alpha_{acc}}$ ) are considered to carry the resistance gene

[ VIDEO 2 ]

https://youtu.be/vcJeKMHbT_g

**Video 2. Simulation of the evolution of a plasmid community considering three incompatibility groups**. See the caption of Video 1 for more details about the visualization.

[ VIDEO 3 ]

https://youtu.be/5DyYBYcr_DU

**Video 3. Simulation of the isolated dynamics of the non-transmissible plasmid maintained in the community from Video 2 comprising three incompatibility groups**. With plasmid costs *α* = 0.01, referring to a 1% reduction of the growth rate compared to plasmid-free cells. See the caption of Video 1 for further details about the visualization.

### Intransitive dynamics stably maintains a set of diverse plasmids in local communities

The composition of the evolving plasmid community can reach a quasi stable state, involving at least one non-transmissible and one conjugative plasmid type per (remaining) incompatibility group (Figure 2c). The evolution of a diverse plasmid community is therefore characterized by the emergence of a network of intransitive substructures (Figure 1b), which enables the maintenance of plasmid diversity itself. If conjugation is considered to be free of cost, plasmid types of the same incompatibility group are not undergoing intransitive dynamics and the evolving plasmid communities are less diverse (see Robustness test on “Cost of conjugation” below).

### Random selection of competitive plasmids causes differences in the plasmid content of local communities

To test the generality of the observed pattern, we made a series of repetitions and compiled some summarized statistics (Figure 3a), assuming that any of these repeated simulations represents a local community of a fragmented population, e.g. similar to the conditions in soil that can lead to spatial separation of soil pores^52^. These simulations confirm our previous results and show how random plasmid survival in local communities is. Although there are fitness differences between plasmids, plasmid survival is not deterministic. It is influenced by random genetic drift, here referring to changes in plasmid frequency due to random sampling of plasmids that successfully reproduce by vertical or horizontal transmission. Otherwise, it is also influenced by selection, as competition rapidly eliminates plasmid types with the lowest fitness regarding their vertical and horizontal transmission efficacy. Only those plasmid types that represent an optimized trade-off between costs and mobility can persist. Very costly plasmids are maintained, but only by high conjugation rates. Non-transmissible plasmids are only maintained if they have very low costs, since competition effectively eliminates all non-transmissible plasmids of the same incompatibility group that exert higher costs, but do also not provide any beneficial function for the host cell. When more incompatibility groups are present, more costly non-transmissible plasmids are able to persist, which shows that purifying selection appears to be less effective considering a higher degree of interaction. If all plasmids belong to the same incompatibility group, non-transmissible plasmids can survive when conjugative plasmids are present because they infect the plasmid-free cells, which would otherwise displace the cells carrying only non-transmissible plasmids, as it has been previously shown^30^. This mechanism does not work when the conjugative plasmids are not more costly (see ‘Robustness tests’ on the ‘Cost of conjugation’ below). The proportion of plasmid types that are non-transmissible can also be higher when more incompatibility groups are present. This is due to a higher proportion of plasmid-bearing cells and a higher plasmid load of individual cells, which alters the meaning of individual plasmid costs, since the relative effect of some additional plasmid cost decreases.

**Figure 3.**
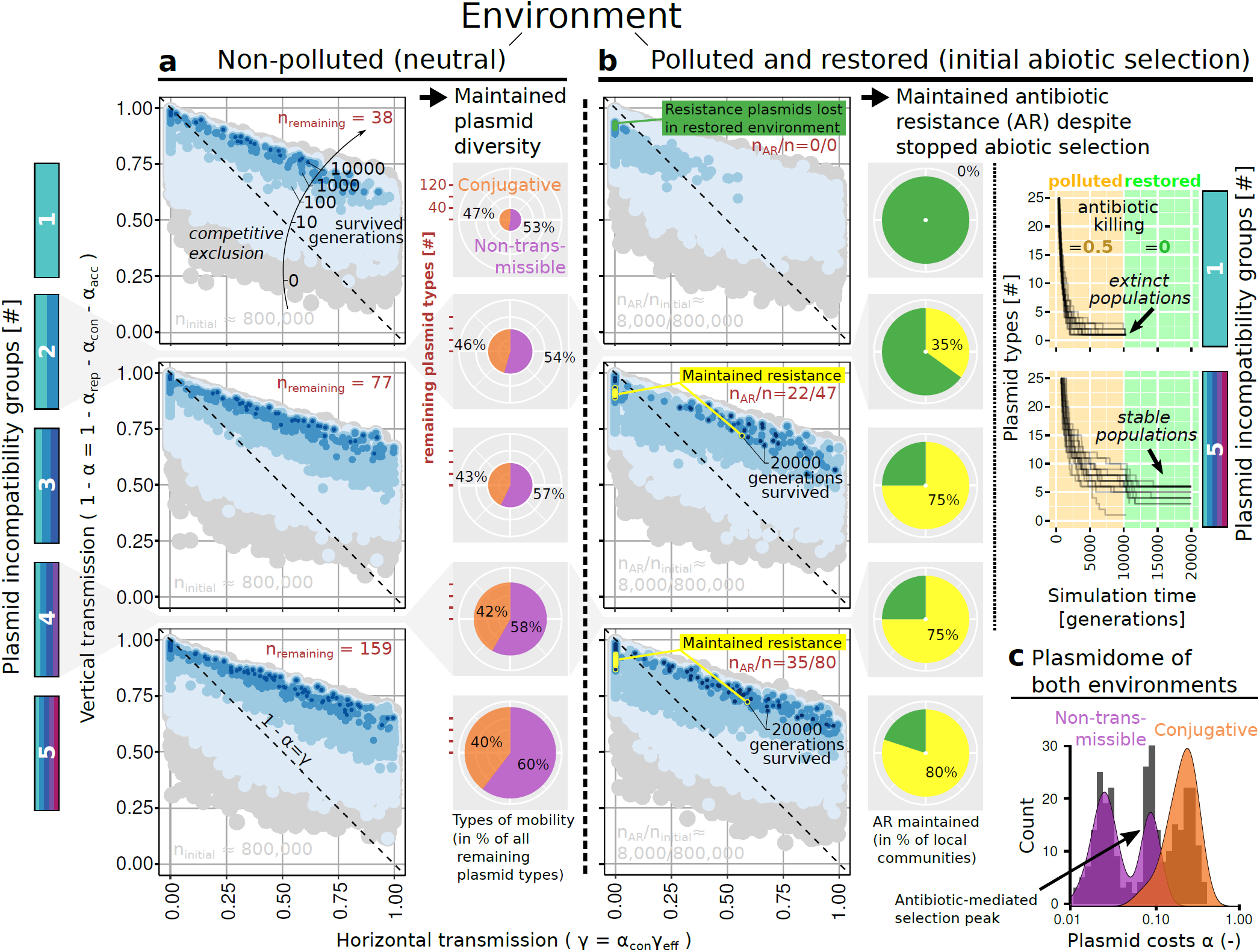
Plasmid content of multiple independently evolved local communities. Each subfigure depicts the outcome of 20 repetitions for the respective setting of environment (neutral vs. initially contaminated with antibiotics) and number of incompatibility groups (1, 3, 5 in detail as well as 2 and 4 in small subfigures). Plasmid diversity increases with the number of incompatibility groups. Plasmid interactions in communities comprising multiple incompatibility groups maintain (**a**) a high plasmid diversity in the absence of abiotic selection of plasmid-encoded functions and (**b**) plasmids conferring costly antibiotic resistance (AR) even though selection by an antibiotic that kills sensitive cells is stopped (here after 10000 generations). **c**: Association between costs and mobility of all plasmid types persisting in one of the presented scenarios (‘non-polluted’ and ‘polluted and restored’; see Figure 3-figure supplement 1 for more details). *n*_*initial*_ … the total number of diverse plasmids in all 20 repetitions of a respective setting; *n*_*remaining*_ … the number of plasmids that survived (shortly given as *n* in (**b**)); *n*_*AR*_ … number of plasmids conferring resistance to antibiotics; Grey: at the beginning of a simulation; Blue colored area: plasmid content after 10, 100, 1000, 10000 (and 20000) generations; Dark red: numbers at the end of a simulation (10000 (**a**) or 20000 (**b**) generations).

### Plasmid interactions maintain antibiotic resistance despite stopped abiotic selection

To investigate the associated effects of antibiotic-mediated plasmid selection, we extended the model considering that an antibiotic is constantly present and kills sensitive cells with a certain probability *ν*, but some plasmids confer resistance to this antibiotic. Our results show that plasmid diversity can be maintained in the presence of antibiotics if the present plasmid types belong to more than one incompatibility group (Figure 3b). Moreover, even when abiotic selection by antibiotics stops, plasmids encoding costly antibiotic resistance can persist. If there is only one incompatibility group, plasmid diversity is more easily lost, since only one plasmid conferring resistance at the lowest costs could remain from the time of abiotic selection, e.g. non-transmissible plasmids. As a result, no other plasmids are present that could help to preserve the costly plasmid providing antibiotic resistance, when this is no longer beneficial. In contrast to this, they can be maintained in plasmid communities composed of more than one incompatibility group. These can exhibit similar temporal dynamics as plasmid communities that evolved in non-polluted environments (Figure 2-figure supplement 1 and Video 4). In fact, the more diverse the evolved communities are, the more likely it is that antibiotic-resistant plasmids will survive.

[ VIDEO 4 ]

https://youtu.be/0mYz1rDW0r8

**Video 4. Simulation of the evolution of a plasmid community under the presence of antibiotics considering three incompatibility groups and varying plasmids providing antibiotic resistance to their hosts**. The simulation is based on a default parameterization, starting with an initially hyperdiverse and completely mixed plasmid community whose progression is followed over 20,000 generations, of which the first 10,000 an antibiotic killing rate of 50% of sensitive cells is present in addition to a “natural” mortality or dilution probability of 20%. When a bacterium and its plasmids are removed from the model domain, a black colored empty space is left that can be recolonized by neighboring bacteria according to their fitness that can be affected by a certain plasmid load (colored plasmids representing (varying) non-transmissible and conjugative plasmids of certain incompatibility groups versus white plasmid-free cells). Please note the monitor that shows the remaining number of diverse plasmids and the ‘tp-pb-relation’-plot depicting their horizontal and vertical transmission fitness similar to Figure 2c. Although the bacteriostatic antibiotic action (‘baa’) drops to 0 after 10,000 generations, one of the plasmid types conferring costly antibiotic resistance (AR, yellow colored) is maintained in the evolved community.

In both simulated types of environments, ‘non-polluted’ and ‘polluted and restored’, the remaining plasmids survived more than 10,000 generations without environmental selection for plasmid-encoded traits. Interestingly, an evolution of plasmid communities under environmental selection for plasmid-encoded traits enabled more costly plasmids to survive, not only over the period of selection, but also in the long term (Figures 3-figure supplement 1, 3a and 3b). This suggests a long-term effect that preserves those genes that were useful at least until recently, which is beneficial for the survival of bacteria under variable environmental conditions. It also indicates, however, that the removal of antibiotics does not necessarily eliminate resistance.

### Robustness tests

To check how strong our results depend on the underlying model assumptions, we applied a series of robustness tests, which is essential for the understanding of the model itself and the identification of ‘robust theories’ about the functioning of the ecological system^53^. In other words, there is no single “true” description of a phenomenon in complex systems, so it is more informative to handle several at once^50^. This is the objective of the following robustness tests.

### Isolated dynamics of pairs of incompatible plasmids selected by plasmid community evolution

The evolved plasmid communities comprise varying combinations of non-transmissible and conjugative plasmid types that belong to the same incompatibility group. Although the previous results demonstrated that these are able to survive in the long term when embedded in their evolved communities, it is not clear whether the dynamics of the evolved communities depends on evolution itself, since this is also associated with spatial self-organisation in our model. We have therefore tested whether the same communities or parts thereof can become established in a population of plasmid-free bacteria. Using the example of the evolved community previously shown in Video 2, it was found that all but the non-transmissible plasmid of incompatibility group 1 can survive if all plasmids of the evolved community are embedded in isolation in an initially plasmid-free population. The non-transmissible plasmid from Inc. group 1 is also not able to survive alone, as it was shown before in Video 3. Considering the plasmids from the other two incompatibility groups of this evolved community in separation, it was found that the inc. 2 plasmids do not survive in isolation (Video 6) while the inc. 3 plasmids do (Video 7). It was also found that the set of plasmids that evolved in the absence of diverse incompatibility groups - as shown in Video 1 - is able to become established in a population of plasmid-free bacteria (Video 8). This suggests that some plasmid communities may only be able to survive if they have undergone spatial self-organisation. Moreover, this seems to be of greater importance for complex communities that include several incompatibility groups than for less complex ones.

[ VIDEO 5 ]

https://youtu.be/sN94ZkgsNLI

**Video 5. Simulation of the dynamics of the plasmid community from Video 2 that evolved considering three incompatibility groups**. Inc. group 1: Non-transmissible plasmid *N* with costs *α*_*N*1_ = 0.012, referring to a 1% reduction of the growth rate compared to plasmid-free cells; Inc. group 2: *α*_*N*2_ = 0.020, conjugative plasmid *C* with plasmid costs *α*_*C*2_ = 0.20 and transfer probability *γ*_*C*2_ = 0.57; Inc. group 3: *α*_*N*3_ = 0.026, *α*_*C*3_ = 0.37 and *γ*_*C*3_ = 0.95. At the beginning, cluster with a radius of 10 cells width are generated next to each other for each plasmid type. See the caption of Video 1 for further details about the visualization.

[ VIDEO 6 ]

https://youtu.be/_FTXWUkFrzc

**Video 6. Simulation of the dynamics of an isolated set of incompatible plasmids that survived during evolution of a plasmid community considering *three* incompatibility groups** (Video 2). Here, inc. group 2: Non-transmissible plasmid *N* with plasmid costs *α*_*N*2_ = 0.020 and conjugative plasmid *C* with *α*_*C*2_ = 0.20 and conjugation probability *γ*_*C*2_ = 0.57. Although both plasmids could survive together in the presence of additional plasmids from at least one other incompatibility group (Videos 2 and 5), they cannot successfully colonize a population of plasmid-free bacteria on their own, because the conjugative plasmid gets into a spatial separation without access to clusters of plasmid-free cells and dies out due to strong competition of surrounding cells with a less costly incompatible plasmid.

[ VIDEO 7 ]

https://youtu.be/YnvgdTPu6kY

**Video 7. Simulation of the dynamics of an isolated set of incompatible plasmids that survived during evolution of a plasmid community considering *three* incompatibility groups** (Video 2). Here, inc. group 3: Non-transmissible plasmid *N* with plasmid costs *α*_*N*3_ = 0.026 and conjugative plasmid *C* with *α*_*C*3_ = 0.37 and conjugation probability *γ*_*C*3_ = 0.95. In contrast to the pair of plasmids from the other inc. group (Video 6), these plasmids successfully co-colonize a plasmid-free population. Although the conjugation probability of the conjugative plasmid is exceptionally high and it would be capable of infecting the entire population, it has a low abundance because it is rapidly displaced by fast-growing cells carrying only the incompatible, non-transmissible plasmid type.

[ VIDEO 8 ]

https://youtu.be/CKUxbJlKjgg

**Video 8. Simulation of the dynamics of an isolated set of incompatible plasmids that survived during evolution of a plasmid community considering *one* incompatibility group (Video 1)**. Non-transmissible plasmid N has some plasmid costs *α*_*N*_ = 0.02 and conjugative plasmid has some plasmid costs *α*_*C*_ = 0.13 and a conjugation probability *γ*_*C*_ = 0.40.

### Absence of plasmid diversity - pairwise plasmid-host interactions

As a validation of the general model behavior, model runs were performed considering only a single plasmid type. It was found that, in the absence of plasmid diversity, sufficiently infectious conjugative plasmids can persist, but non-transmissible plasmids not (Figure 4), which is in line with the predictions of previous studies assuming similar conditions (neglected co-occurrence of diverse plasmid types), as summarized in the introduction. In addition to Video 3 that showed the spatio-temporal dynamics of a single non-transmissible plasmid and its continuous decrease until extinction, the dynamics of a single conjugative plasmid that rapidly infects the whole population of plasmid-free cells is shown in Video 9. Figure 4 further indicates that plasmid frequencies might also settle around intermediate frequencies. This is shown and explained with Video 10.

**Figure 4.**
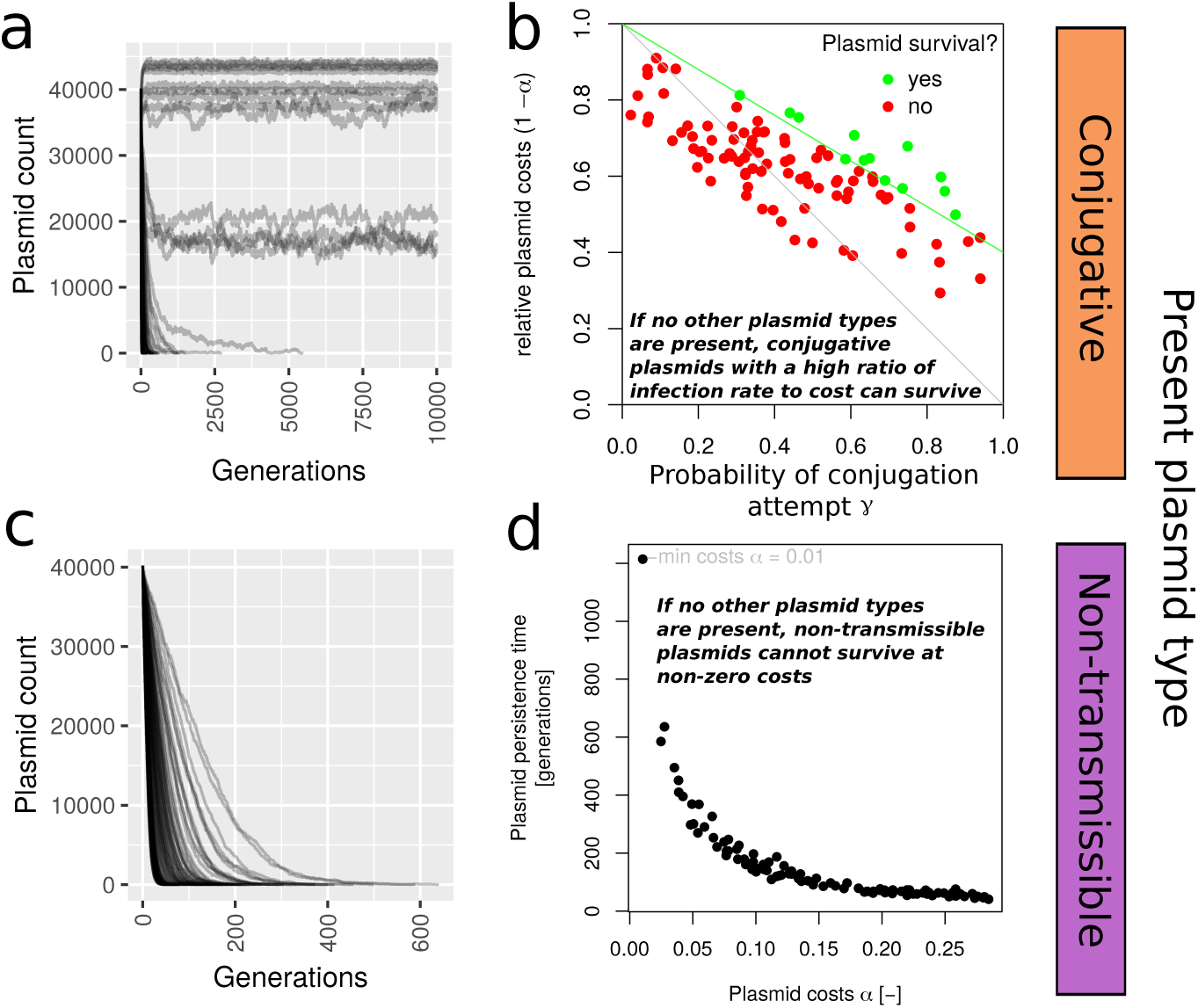
Plasmid persistence in the absence of plasmid diversity. Considering the presence of either one single conjugative or one non-transmissible plasmid type, conjugative plasmids are able to survive, when the probability to perform a conjugation attempt is far greater than the associated plasmid costs (a, b). As a result, tested plasmid types most often either got lost or were highly abundant. The dynamics of some plasmids show fluctuations due to spatial effects caused by the formation of clusters of plasmid-free and plasmid-bearing cells (Lines represent conjugative plasmid counts only). Non-transmissible plasmids are not able to survive in the long-term in a non-diverse plasmid community, given that their costs are greater than zero and they do not provide beneficial functions for their hosts (c, d); Lines (a, c) represent independent simulation runs that are also represented by points (b, d).

[ VIDEO 9 ]

https://youtu.be/4M5nQuhaW8g

**Video 9. Simulation of the dynamics of a single conjugative plasmid**. Here, the conjugative plasmid that belongs to the pair of inc. group 2 plasmids that remained as part of the evolved community shown in VIDEO 2 is used. It has plasmid costs *α* = 0.37 and a conjugation probability *γ* = 0.95. The plasmid rapidly infects nearly all plasmid-free cells and is able to survive in the long term on its own. Note that the same plasmid got extinct due to spatial separation in another setting with hierarchical competition of a co-colonizing, incompatible non-transmissible plasmid with lower costs (Video 6).

[ VIDEO 10 ]

https://youtu.be/gp9Y-5Hodmc

**Video 10. Simulation of the dynamics of a single conjugative plasmid settling at intermediate frequencies in a population**. Here, the infection dynamics of a conjugative plasmid with costs *α* = 0.3 and a conjugation probability *γ* = 0.5 is examined. Although the initial increase in frequency is very steep, the plasmid is not able to infect the whole population. This is because the probability of finding a potential recipient in the neighborhood decreases the more cells are infected. When more than half of the cells carry plasmids, the spatial pattern is inverted, i.e. it undergoes a transition from a pattern representing a number of clusters of plasmid-carrying cells in a pool of plasmid-free cells to the opposite. As a result, there are on average more plasmid-carrying cells than plasmid-free cells at the border between the two aggregates, which reduces the success rate of horizontal gene transfer and can lead to a fluctuation around this state.

### Population size and more than five incompatibility groups

It was tested if the evolution of a stable state that maintains a certain number of diverse plasmids is affected by the model world dimension and a larger number of incompatibility groups. This test revealed that the evolving local communities are able to preserve a larger number of diverse plasmid types, the larger the model world size and the number of incompatibility groups (Figure 5). Smaller populations are only able to sustain a smaller diversity of both plasmid types and incompatibility groups. The increase of plasmid diversity with the number of incompatibility groups is limited by the size of the local population. These results are consistent with previous studies indicating that effective population size has a strong influence on pangenome size^3^.

**Figure 5.**
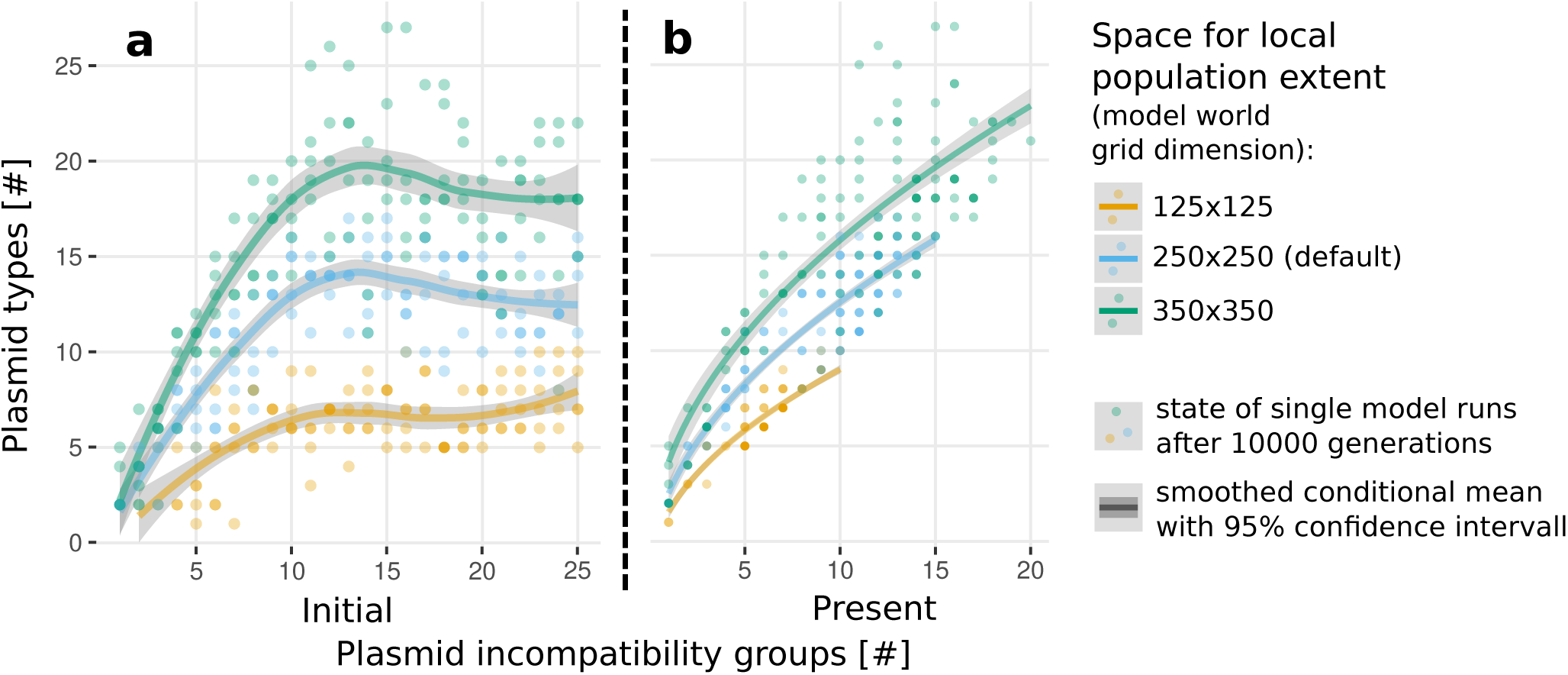
Plasmid diversity in evolved communities considering varying population sizes and numbers of incompatibility groups. The initial number of plasmid incompatibility groups is determined by initialization, whereas the present number reflects the state of each local community (single model run) after 10000 generations.

### Simulation time

To test if the community composition that emerged after 10000 generations significantly changes in an increased time horizon, we ran a couple of simulations for 30000 generations. The results show that the number of surviving plasmid types does not further or not substantially change in an increased time horizon (Figure 6). Along with these simulations a model version considering another mechanism of incompatibility was tested, as our next robustness test shows.

**Figure 6.**
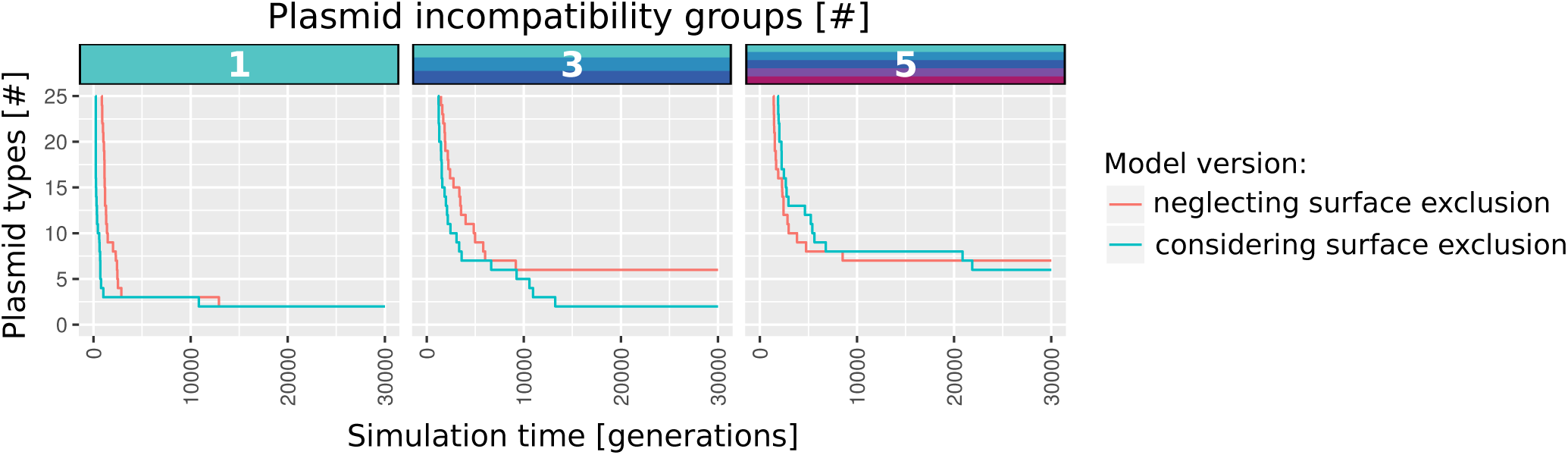
Model behaviour considering varying mechanisms of incompatibility. Each line depicts plasmid diversity dynamics for a single model run either assuming surface exclusion of incompatible plasmids (model’s default) or random exclusion during cell division, when incompatible plasmids co-occur in the same cell. Neglecting the effect of surface exclusion, more plasmids can enter cells, enhancing plasmid maintenance and the stability of the evolved community. In reality, some plasmids actively promote surface exclusion whereas others don’t. But as both mechanisms alone show a similar behavior, this appears to be unimportant for our main findings.

### Incompatibility mechanism

In the standard model version it is assumed that incompatibility prevents the entry of plasmids into a cell that already contains another plasmid of the same incompatibility group. We tested this simplifying assumption and performed simulations with a model version that neglects such a surface-exclusion mechanism. Instead, incompatible plasmids might co-occur in the same cell, but are easily lost during cell division. We included this feature in our model using probabilities to generate daughter cells that are free of one of both incompatible plasmid types, when a cell harboring more than one incompatible plasmid type performs cell division. This imitates that replication is usually disrupted in such cases, resulting in decreased copy numbers and increased segregation rates. Incompatible plasmids are thus also excluded from the same cell in this way, only with a slight delay in which horizontal gene transfer can potentially take place. Our test results show that this alternative model definition does not seriously affect the model behaviour (Figure 6). This enabled us to opt for the computationally less demanding mechanism of surface exclusion.

### Immigration

To test how the immigration of bacteria that bear new plasmid types influences the stability of the evolved community, we extended the model by a subroutine for immigration. Within this submodel, a random lattice cell can be occupied by an immigrating bacterium with a probability *ϖ*, for the presented test cases fixed at 10^−5^ per lattice cell per time step (corresponds to one hour). Immigrants can be plasmid-free or plasmid-bearing according to the same probabilities that were used to set up the initial state of the model (Table 2). First, it was tested if the invasion of diverse plasmid types affects the stability of the evolved local community. Second, it was tested if the same plasmid community evolves when the model is initialized with a single plasmid type rather than a diverse community, but considering that new plasmid types can immigrate with bacteria. Figure 7 shows exemplary plasmid community dynamics for both above mentioned cases. These results confirm that plasmid diversity can be maintained for evolved communities and evolves in initially clonal populations considering immigration. Both the stability of the proposed plasmid diversity mechanism and the behavior of the more simplified model version that neglects immigration is therefore robust, since the resulting plasmid communities are similar.

**Figure 7.**
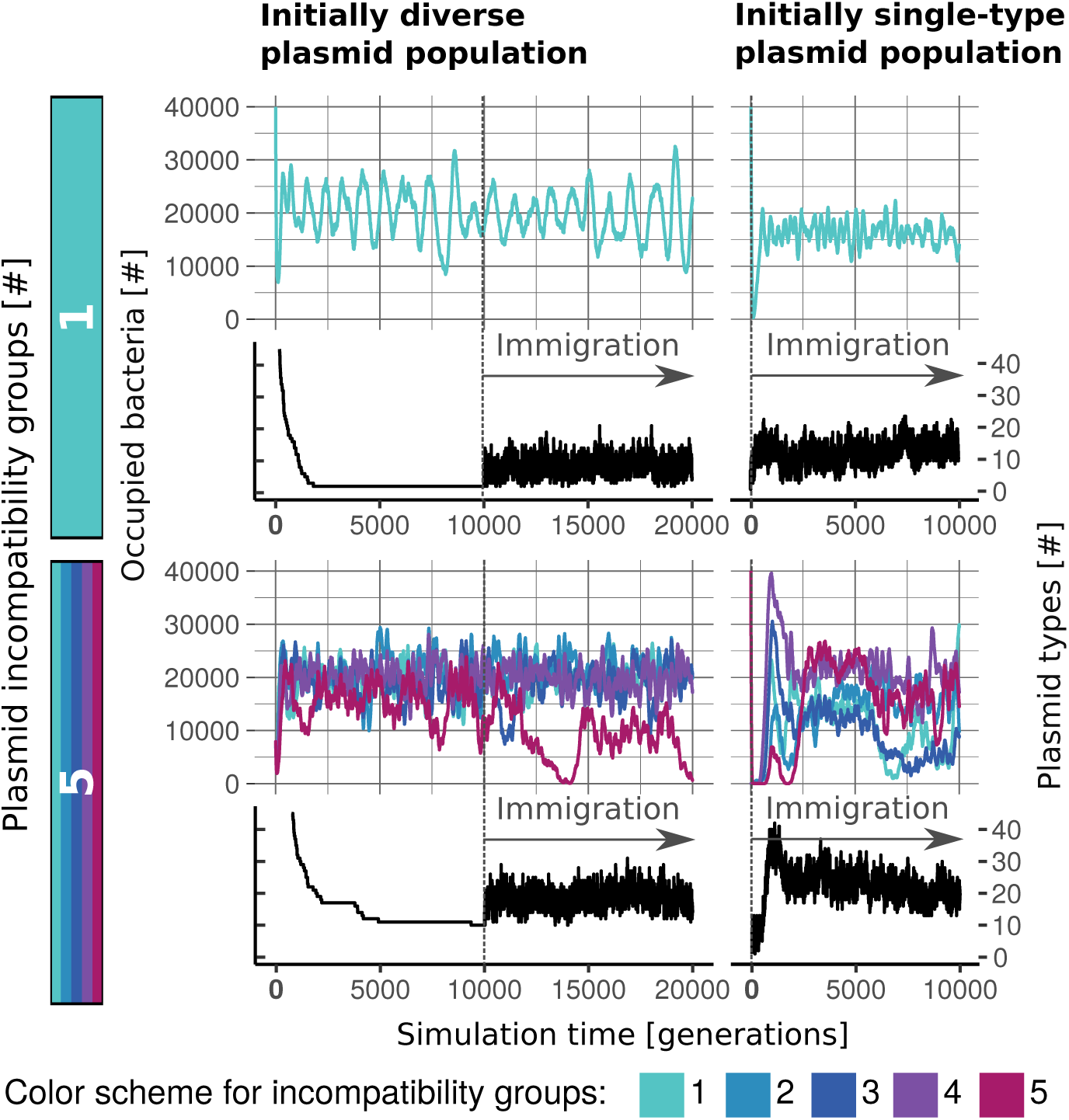
Plasmid diversity can be maintained for evolved communities and evolves in initially clonal populations considering immigration. Each row represents the observations for a single simulation run assuming initially 1 or 5 incompatibility groups and a probability to become occupied by an immigrant of 10^−5^ per lattice cell per hour. Immigrants are random plasmid-free or plasmid-bearing bacteria. Although plasmid diversity is constantly raised by a kind of noise resulting from immigration, this does not substantially affect the frequencies of the prevalent plasmid types.

### Well-mixed, unstructured environments

Another robustness test focused on local interaction, i.e. the interaction with neighboring bacteria, and spatial structure. To observe if plasmid diversity can be maintained for the same reasons in a mixed population, we extended the model by a subroutine for mixing. This procedure relocates any cell after each time step, which prevents the emergence of a spatially clustered community structure. Even in this case, plasmid diversity was maintained, but antibiotic resistance got lost in restored environments (Figure 8). The presence of a structured environment such as biofilms can therefore be seen as an important factor that supports the persistence of costly traits such as antibiotic resistance, which might get lost in a well-mixed environment such as the planktonic phase of water bodies.

**Figure 8.**
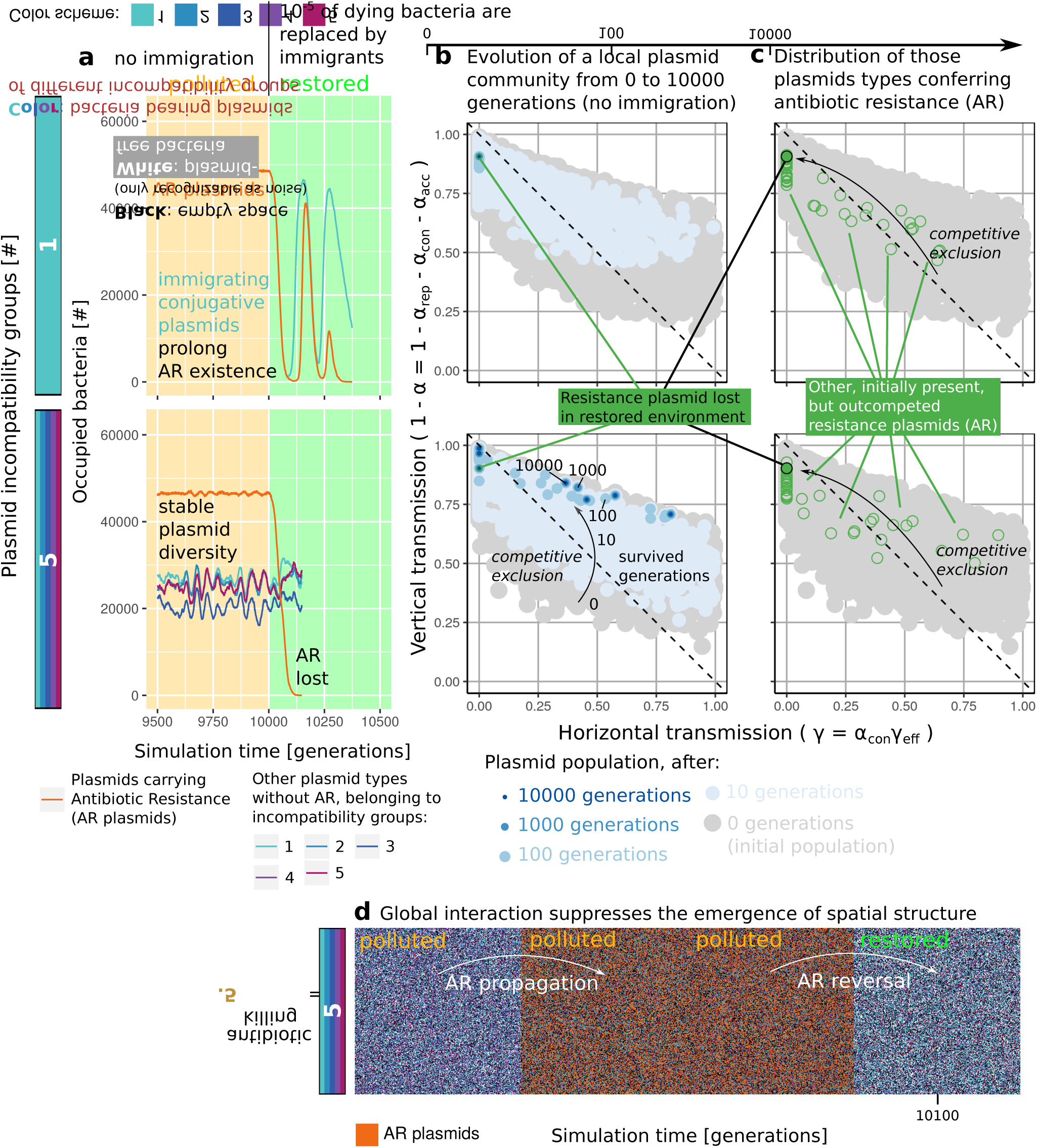
Plasmid persistence considering environmental mixing. **a**: Plasmid diversity can be maintained in polluted environments when more than one incompatibility group is present; Antibiotic resistance (AR) is lost in restored environments, even considering immigration that could help to reestablish a stabilizing community structure; **b**: Depending on the number of plasmid incompatibility groups, only some of the most competitive plasmids can be sustained in each local plasmid community; **c**: Although multiple plasmid types provide AR, competitive exclusion reduces such redundancy by a selection of one of the most competitive plasmid types that confer this function to the local community; **d**: Constant mixing of bacteria prevents an evolving community to become spatially structured

### Distribution of antibiotic resistance among plasmid types

Antibiotic resistances cannot persist after stopped antibiotic use when they are conferred by a single non-transmissible or weakly to moderate infectious conjugative plasmid (Figure 9). This is because such systems are hierarchical during the timecourse of antibiotic use, since all other plasmids or at least those of the same incompatibility group that do not confer the antibiotic resistance gene are likely eliminated. Therefore, the conditions of the non-hierarchical tournament cannot be fulfilled, when antibiotic-mediated selection stops, which facilitates resistance reversal. Conjugative plasmids that are infectious enough to compensate for the huge costs imposed by the non-beneficial resistance might still persist, even in the absence of abiotic selection. However, this only works as long as it does not have to cope with other less costly plasmid types of the same incompatibility group. Otherwise, the requirements for the non-hierarchical tournament maintaining the resistance can also be established, when a plasmid mutant arises that confers the resistance, but lost or at least repressed its conjugation genes, making this mutant less costly than the other conjugative type that is still present. The investigation of such conditions requires a consideration of transiently derepression, which has been previously approached using a similar model^30^. Since such adaptations are likely, we argue that the maintenance of antibiotic resistance by single plasmid types is probably less restricted than these tests suggest.

**Figure 9.**
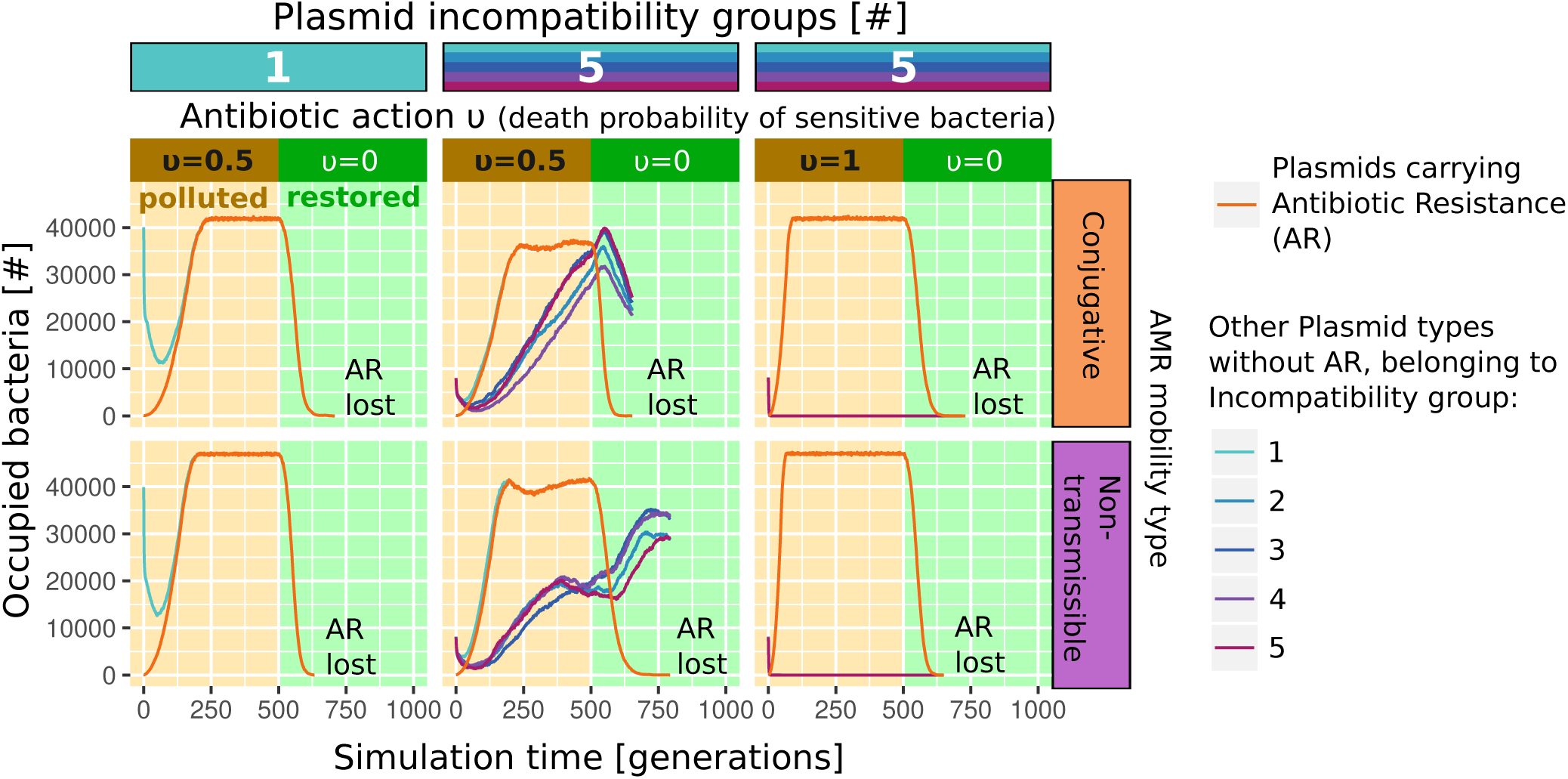
Resistances are lost considering single plasmid types conferring antibiotic resistance genes. Each plot depicts the results of a single simulation, initialized with an antibiotic resistance plasmid (ARP) that has the mean properties of the initial plasmid community (see table 2 for a list with means for plasmid module costs, efficiency of conjugation etc.)

### Cost of conjugation

The model that our main results are based on considers that plasmids with high conjugation rates are usually more costly, as we defined the conjugation probability to depend on the cost of the conjugation module *α*_*con*_ and the conjugation efficiency *γ*_*e f f*_, which represents a factor around 2 *±* 1 (see Table 2) for the translation of costs to horizontal transmission fitness *γ*. Although we generally assume that conjugation is costly for the individual host cell, a significant portion of the population may in fact suppress plasmid transfer functions^54^, which reduces the plasmids cost-of-carriage for these cells. To test whether the assumption that conjugation is costly seriously influences our general results, we performed simulations of ‘non-polluted’ (Figure 10) and ‘polluted and restored’ environments (Figure 10-figure supplement 1) in which conjugative plasmids do not have additional costs (see Figure 1-figure supplement 1 for the composition of traits in the initial plasmid community). These show that plasmid diversity is still maintained, but nearly all remaining plasmids are conjugative and have very low costs. For 3 and 5 incompatibility groups, essentially no plasmid-free cells are left. This composition and the associated dynamics of individual plasmid types (Figure 10-figure supplement 2) shows that no intransitive system emerges if conjugation is free of costs. In this case, diversity is maintained by plasmid incompatibility, spatial dynamics and other stochastic effects that allow inferior competitors (e.g. less infectious conjugative plasmids) to escape from deterministic competitive exclusion. Non-transmissible plasmids only have a chance of survival if all conjugative plasmids of the same incompatibility group are lost. In polluted (and restored) environments, the relative importance of stochastic effects is decreased due to selection of plasmid-encoded traits such as antibiotic resistance. This is similar to the behaviour of our default model version. However, we conclude that the default model version showed a better agreement with empirical observations because it helps to explain how non-transmissible plasmids, which account for about half of the known plasmids^55^, can survive. It is also consistent with observations suggesting that a significant proportion of bacteria are plasmid-free^56^.

**Figure 10.**
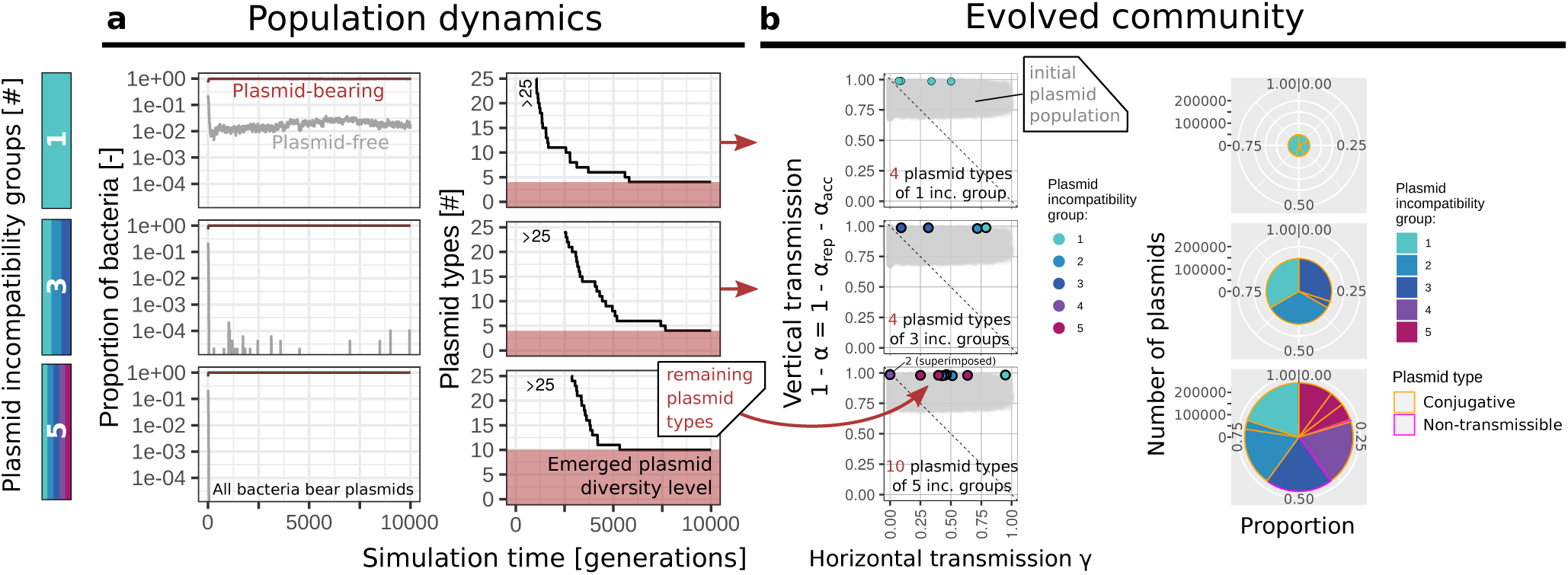
Assuming that there is no costs for conjugation, diverse plasmids are maintained in a non-polluted environment (no abiotic selection) mainly by neutral interactions due to nearly non-existent host fitness differences and a decreased importance of conjugation. Each row depicts the results of a single simulation, initialized with a plasmid community consisting either of 1, 3 or 5 different plasmid incompatibility groups. The frequency of plasmid-free cells is or approaches 0 and for each incompatibility group either conjugative or non-transmissible plasmid types survive. The composition of the evolved plasmid communities indicates that diversity is not maintained due to rock-paper-scissor dynamics, but that in principle inferior competitors such as less infectious conjugative plasmids can escape from deterministic competitive exclusion due to a generally weak importance of conjugation when all plasmids have a high vertical transmission fitness and nearly no recipients (e.g. plasmid-free cells) are left.

## Discussion

Natural communities that generate subpopulations with varying plasmid content are at an advantage, as genetic variation is the key for bacteria to cope with environmental uncertainty^57^. It has been shown that conjugation rates of diverse plasmids from varying incompatibility groups are sufficiently high to maintain them in the absence of positive selection for the plasmid encoded trait^33^. Previous findings^30^ demonstrated that this property can also facilitate the persistence of other, less infectious plasmids by intransitive dynamics. The results presented here suggest that such intransitive dynamics of plasmids with different mobility is an emergent property of plasmidomes and could therefore be an important mechanism for the preservation of genetic diversity in nature.

By means of an individual-based model of the plasmidome, we observed the evolution of a population consisting of multiple plasmid types that differ in cost, transfer probability and incompatibility and that interact inter- and intra-cellularly with each other. We found that these biotic interactions between diverse plasmid types lead to the evolution of a system lacking strict competitive hierarchies between plasmids with different mobility traits. We did not consider explicit evolutionary changes at the gene level, but we observed how dynamic rearrangements of the plasmid composition on the bacterial level alter the competition between various plasmid-host associations. This results in a niche differentiation of plasmids that enables the preservation of plasmid diversity and the persistence of costly plasmid-encoded traits in the absence of abiotic selection. When the evolved plasmid community spans multiple incompatibility groups, the interactions between plasmids represent a complex network that can provide niches even for plasmid types such as non-transmissible plasmids conferring costly antibiotic resistance that could otherwise not survive. Such intransitive competition has long been overlooked, but is now recognized to shape species coexistence for a variety of taxa at various ecological scales^58^. It has for example been shown that the interaction between antibiotic production and degradation enables coexistence in microbial communities^59^. Our results also fit well with the observation that interactions in more complex communities may limit the response to abiotic selection, as recently demonstrated for competitive bacterial species^60^ and across trophic levels^46^.

The revolution in sequencing methods^40^ provided cultivation-independent approaches that can be used to study entire plasmid pools of environmental samples^61^ as well as to link resistance genes to plasmids and microbial taxa^62^. Plasmidome analysis have been carried out, amongst others, in freshwater^7^, soil^38^, wastewater^35,39^, groundwater^44^ as well as in the rat recum^37^ and the bovine rumen^36^. This revealed, for example, differences in the composition of replication families between wastewater and soil plasmidomes^38^. Single studies also showed that, for example, a groundwater plasmidome can contain hundreds of plasmids that span multiple incompatibility groups^44^, which is a finding consistent with our general modelling results. Some abundant plasmids were found to encode traits such as metal, antibiotic and phage resistance along with toxin-antitoxin systems^44^. This is a feature that we did not considered in the model, but that is likely enhancing the survival of plasmids that are much more costly or less infectious. The content in accessory functions of plasmidomes likely differs between environments that exert different abiotic pressure. It has for example been found that the plasmidome of biological wastewater treatment reactors supplemented with antibiotics contains a multitude of antibiotic resistance genes^39^, whereas some environments such as freshwater have been shown to contain a high diversity of plasmids with no or unknown accessory functions^7^. This fits well to our simulation results for polluted and non-polluted environments: multiple plasmids providing certain costly accessory functions can persist in polluted (and restored) environments, but are more likely to go extinct in plasmid communities evolving in non-polluted (pristine) environments.

Horizontal gene transfer can be considered to be of major importance for the exchange of genes within and between environmental microbiota^63^. Whereas some plasmids are in reality limited in their host range, others have been found to frequently perform even inter-Gram transfer^64^. The distinction between the narrow-host-range and broad-host-range is not considered by our model as it only considers a clonal bacterial population, but we consider this to be an important aspect for future research. For example, some bacteria such as *Aeromonadaceae* have been identified to be major reservoirs of antibiotic resistance in wastewater and to carry IncQ plasmids that were observed to have the broadest host ranges among plasmids^62^. Such combinations of reservoir and vector likely enhance the spread of antibiotic resistance to other members of the microbial community.

Physical fragmentation of natural habitats such as soils has been demonstrated to affect the transfer of genetic information in support of microbial diversity^52^. If we similarly consider that repeated simulations of our model represent a natural community consisting of fragments that are limited in size and degree of interaction, we find that the stochastic composition of plasmid communities evolving on a local scale increases the diversity on the population level. We suggest this stochasticity to be of high importance for the preservation of an increased genetic diversity. We consider this behavior to be realistic from a theoretical point of view, because the survival of a particular plasmid is probably not deterministic, given that many other plasmid types with similar fitness characterisitics are likely present, evolve or immigrate and compete to survive in a dynamic plasmid community.

The properties of the plasmids that are able to survive as part of the evolved communities in our simulation experiments (Figure 3c) show similarities to patterns reported from bioinformatic analysis of plasmids in the GenBank database^55,65^: a multimodality of the plasmid size distribution that can be best interpreted when plasmids are divided into classes such as non-transmissible and conjugative. In our model plasmids are composed of gene modules that are associated with costs. Such a direct cost to size relationship might not hold considering that small plasmids are often present in more copies per cell. But it holds if we consider additional genes to be more disruptive or demanding for the host cell than copies of some already existing genes. If, for example, a plasmid provides antibiotic resistance, these additional genes increase plasmid size as well as plasmid costs, which has been observed by fitness increases after deletions of plasmid-borne antibiotic resistance genes^66^. Additional resources are also required for the expression of other additional functions such as the conjugation machinery but can be alleviated either due to the silencing of conjugation genes^54^ or their complete deletion, which has been reported for some plasmids that accordingly have a smaller plasmid size than their ancestors^65^. Similarly, non-transmissible plasmids in our model can be seen as a transfer-deficient variant of conjugative plasmids. Non-transmissible plasmids of larger costs (and size) were able to survive in our simulations, when their increased costs were at least beneficial during the evolution of the plasmid community (e.g. long-term persistence of plasmids encoding antibiotic resistance, when the environment was initially contaminated with antibiotics). This suggests that abiotic selection for plasmid encoded traits may have an effect on the pangenome that lasts longer than its direct occurrence and could also influence patterns of plasmid size and mobility. Such system-immanent stability of genes is presumably an advantage, since a lot of accessory functions might only rarely be beneficial, but must be preserved until the next drastic change in the environmental conditions under which these genes are required. Mechanisms that support the preservation of genes in the common gene pool are therefore likely favored by natural selection, which might help to explain why plasmids varying in mobility and incompatibility evolved.

In summary, we conclude that established interactions between diverse plasmids in evolved plasmid communities are likely to preserve a huge pool of genetic information that is highly accessible for the microbial community. When abiotic selection of a costly trait such as antibiotic resistance is stopped, interactions among plasmids that span multiple incompatibility groups have been shown to facilitate the persistence of such accessory functions in plasmidomes. We call for studies that further investigate the role of biotic interactions, involving varying plasmid types, bacterial species and other mobile genetic elements, as such complex setups will help us to unravel the impact of such natural characteristics.

## Methods

An individual-based model (IBM) was developed, as this allows to account for individual and spatial variability and to investigate emergent interactions in complex communities. These features are likely important for the life of plasmids in natural microbial communities. Such a bottom-up approach can provide insights that cannot be obtained with classic population level models^67^.

### Model description

Here we present a model description following the Overview, Design concepts and Details (ODD) standard protocol for individual- and agent-based models^68^. It is proposed to facilitate evaluation and comparison of IBM and consists of seven elements, which will in turn be used to address general and specific aspects. The model was implemented using the software platform NetLogo 6.0.2^69^ that works on a Java virtual machine.

#### Purpose

The mechanisms enabling the diversity of plasmids observed in nature are not fully understood. Although plasmids can be beneficial if they encode for any environmentally selected function, they may otherwise represent a burden to their host cell and can become eliminated within a few hundred generations. Frequent environmental selection can maintain single, sufficiently beneficial plasmids, but this mechanism does not explain the diversity of plasmids in size and function as observed in nature. The model primarily addresses whether intrinsic mechanisms, i.e. those that originate from plasmid diversity itself, are capable to regulate and maintain it.

#### Entities, state variables and scales

The model composes two entities: lattice cells and plasmids. Each lattice cell is either empty or occupied by a single bacterium. A bacterium may bear a number of plasmid types that reside as individuals on the respective cell.

State variables of lattice cells are location (integer x- and y-coordinates of the lattice point) and occupation (representing a bacterium or an empty lattice cell). The state variables for plasmids are the association with a certain bacterium (=location), and costs of certain gene modules related to replication, conjugation and accessory functions. The variable conjugation efficiency determines how costs of the conjugation module are related to the realized plasmid transfer probability (if conjugation efficiency = 1, then the costs on vertical transmission are theoretically balanced by horizontal transmission; see another study^33^ for a general description and empirical use). If the absence of conjugation costs is modelled, the plasmid transfer probabilities are calculated in the same way, but the costs of the conjugation module are ignored afterwards. The incompatibility group defines those plasmid types that cannot co-occur in the same cell. All state variables are listed in Table 1. During initialization, random states are assigned to the plasmids, which - besides location - do not change over time and depend on distribution parameters that are used exclusively for initializing the model (Table 2 lists the parameters; Figure 1-figure supplement 1 shows the frequency distributions of plasmid traits of an exemplary plasmid community at initialization). Other constant, but global model parameters are mortality, antibiotic action and segregation.

The model world consists of a grid with a dimension of 250 *×* 250 lattice points and periodic (or ‘wrap-around’) boundaries. It represents thus a surface-attached population of maximal 62,500 bacteria. The model world may correspond to one or a half square millimeter of a biofilm surface, assuming an average bacterial cell size of about 1 or 2 *µ*m and distances between bacteria in the same order of magnitude. Each model time step represents one hour. A certain number of time steps is required to generate as much new cells by bacterial cell division as are given for an existing population. This is defined as the time of one generation. Experiments simulating ‘non-polluted environments’ lasted for 10000 generations and for ‘polluted and restored environments’ 20000 generations.

#### Process overview and scheduling

At each time step, lattice cell states are updated sequently in random order, following the models decision rules (Figure 1-figure supplement 2). For each lattice cell update a random lattice cell is chosen (the so called “focal cell”). If the focal lattice cell is not occupied by a bacterium, nothing happens. If it is occupied either mortality, cell division or horizontal gene transfer happens according to the probabilities for these events. The probabilities depend on the particular bacteria-plasmid association characterizing the current state of the focal cell. Whereas mortality is fixed for each bacteria-plasmid association, the probabilities for cell division and transfer depend furthermore on the local resource availability as well as the transfer probabilities and the costs that are associated with all plasmids of the bacterium (see ‘Submodels’ for further details).

If a bacterium dies, the lattice cell becomes empty and all plasmids die too. If a bacterium performs cell division, a random empty lattice cell in the local neighborhood is occupied by a copy of the bacterium and its residing plasmids, whereas plasmids can get lost according to a fixed segregation probability. If plasmid transfer happens, one of the plasmids that reside concurrently on the host is randomly selected according to its relative capability to perform the transfer. It will then attempt to transfer a copy to a random, non-empty lattice cell in its local neighborhood (Figure 1-figure supplement 3 illustrates the spatial context). Transfer is prevented if the target bacterium already contains the identical plasmid type (always considered for technical reasons: a plasmid type represents a set of traits that is either present or absent, whereas plasmid copy numbers are neglected). The same applies for plasmids of the same incompatibility group.

#### Design Concepts

##### Basic principles

The mutual exclusion of plasmids belonging to different incompatibility groups leads to the emergence of an intransitive system, which lacks strict competitive hierarchies. This is similar to models that describe coexistence as the result of the so-called rock-paper-scissors concept (e.g.^51^). Biotic interactions allow niche differentiation that can preserve biodiversity.

##### Emergence

The temporal dynamics, spatial distribution and (long-term) composition of the plasmid community emerges from the intra- and inter-cellular interaction of plasmid types varying in costs, transfer-rates and incompatibility. Each initial plasmid type has a unique combination of features (costs, horizontal transfer probability and incompatibility group) which may or may not be advantageous in competition with other plasmid types or plasmid-free cells. The evolution of interaction within the plasmidome might be understood as follows: different plasmid-plasmid associations compete within the same host species, resulting in a survival of the fittest that represent an optimized strategy regarding plasmid costs and horizontal transmission. Low-cost plasmids survive because they have a small effect on vertical transmission, even in conjunction with other more costly plasmids. Conjugative plasmids could survive as long as they are infectious enough to compensate for the loss they impose on host fitness^17^. Nonetheless, this balance between costs and function is disturbed, when multiple plasmids are present in the same host cell. The model shows that in such complex communities a network of specific interactions can arise. Each member of the evolving community has a more competitive and a less competitive opponent, which has been suggested to be of central importance for microbial diversity^59^. One of the major differences in our model is that this principle of non-hierarchical competition occurs simultaneously for varying incompatibility groups, forming a network of intransitive substructures (Figure 1b).

##### Adaptation

The model does not consider adaptation of individuals, which means that plasmids in our model cannot adapt their costs, conjugation efficiency and incompatibility group influencing their behavior and fitness. Individual bacteria in our model can also not directly adapt their own fitness, as this is influenced by the residing plasmids. However, the community adapts indirectly over time as only some of the plasmid types can survive which, in co-residence with the other remaining compatible plasmid types, do not cause excessive fitness costs on their bacterial hosts.

##### Prediction

Bacteria and plasmids are not able to predict their behavior and thus optimize it in our model.

##### Sensing

Plasmids are able to sense if a bacterium already bears a plasmid of the same incompatibility group in our model. If so, any attempt to perform horizontal transfer to this bacterium fails. Bacteria do not sense other bacteria in their neighborhood, but the number and costs of plasmids residing on themselves.

##### Interaction

In our model, the interaction of plasmid types occurs by inter- and intra-specific competition. The latter is given when neighboring host cells carry the same plasmid type and consequently compete for the same local resources namely empty neighboring lattice cells. Inter-specific competition is given when neighboring host cells carry different plasmid types. In addition, inter-specific competition of compatible plasmid types occurs on an intra-cellular level, since the fitness of the host and thus the probability of cell division and plasmid spread decreases with the number of hosted plasmids.

##### Stochasticity

The initial composition of traits in the plasmids (costs, horizontal transfer rate and incompatibility group) are drawn from random distributions. All processes are handled by probabilities (Figure 1-figure supplement 2). Cell division, for example, depends (1) on the availability of empty cells in the local neighborhood from which one of them is randomly chosen, and (2) on the host-fitness described as a probability which decreases with the costs of all plasmids carried. Such stochasticity leads to demographic noise in the bacterial population.

#### Initialization

The model is initialized by a random assignment of lattice-point states. The default probability that a lattice point becomes occupied by a bacterium corresponds to 1 minus the probability for mortality *ω*, which approaches the expectable population density of such a system. Each plasmid created receives values for its state variables, defining replication module costs *α*_*rep*_, conjugation module costs *α*_*con*_, accessory module costs *α*_*acc*_, efficiency of conjugation *γ*_*e f f*_ and plasmid incompatibility group *η*. These parameters are plasmid type-specific and drawn from a normal or uniform distribution with some constraints on the lower limit (see Table 2). Half of the initial plasmid community is non-transmissible (*α*_*con*_ = 0). The default probability of a bacterium to carry a plasmid is 0.8, which is similar to some reported frequencies of plasmid-bearing strains of 73% in *Lactobacillus helveticus*^6^. This value is high enough to ensure that the initial diversity of plasmid traits is comparable among simulations. Considering the default parameterization, ca. 40,000 different plasmid types were initially generated.

#### Input data

The model does not use input data to represent processes that change over time.

#### Submodels

The following paragraphs describe in detail the single model processes and the calculation of the used probabilities *P*(*cell lysis*), *P*(*cell division*), *P*(*plasmid trans f er*) and *P*(*no action*) (see schedule given in Figure 1-figure supplement 2). For the latter, local conditions, state variables (Table 1) and fixed parameters (Table 2) are taken into account. In general, parameters are chosen in a way that *P*(*cell lysis*) + *P*(*cell division*) + *P*(*horizontal plasmid trans f er*) *<* 1. The probability that nothing happens when a lattice cell is considered for an update in a single time step is given as follows:

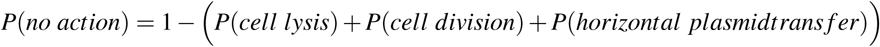

### Cell lysis

A bacterium and all its residing plasmids die according to a global probability for mortality. They are then removed from the model domain, which can occur due to natural mortality but also due to other processes such as washout or predation. In addition, bacteria that do not host a plasmid with the relevant trait can die with a certain probability under antibiotic action. The probability for cell lysis can thus be denoted as:

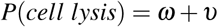

with the global probability for mortality *ω* and the antibiotic action *ν*. Cells that died leave an empty lattice cell that can become recolonized through ‘cell division’ by other bacteria.

### Cell division and vertical plasmid transfer

Plasmids can be inherited within a cell lineage (also called ‘vertical gene transfer’), depending on the growth of their host. A bacterium performs cell division according to a probability that depends on its local resource availability and the costs that are imposed by all residing plasmids. The local resource availability represents a factor that is given by the proportion of empty cells in the 8 cell Moore neighbourhood. Thus, the less neighboring cells are occupied the higher the nutrient availability for the focal cell. The costs of plasmid carriage is given as the sum of costs that are imposed by each residing plasmid. Let us denote the factor given by the local resource availability as *f*_*res*_ and the factor given by the costs of plasmid carriage as 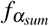, than the probability for cell division follows:

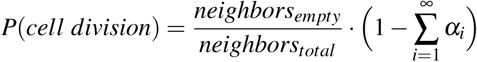

whereas *α*_*i*_ represents the costs imposed by the *i*-th plasmid on the host cells growth rate; *neighbors*_*total*_ refers to the surrounding cells (=8), and *neighbors*_*empty*_ represent the number of empty cells among them.

Considering the exemplary conditions for the focal lattice cell depicted in Figure 1-figure supplement 3, *neighbors*_*empty*_ = 3,c *α*_*i*=1_ = 0.21, *α*_*i*=2_ = 0.02, resulting in *P*(*cell division*) = (3*/*8)(1 − 0.21 − 0.02) ≈ 0.29.

Plasmid heritage during cell division is controlled by two subroutines: *Plasmid incompatibility* and *Segregation*.

#### Plasmid incompatibility

In the standard model version, surface exclusion is considered. This means that plasmid incompatibility already has an effect during the horizontal gene transfer and prevents the co-occurrence of incompatible plasmids in the same cell. Such exclusion systems maintain plasmid exclusivity in a host and can also function via restriction-modification systems or CRISPR-Cas that act on the plasmid upon entry^70^. Entry exclusion systems may not be incompatibility-group specific, as it has been demonstrated for instance for the IncA and IncC plasmids that both also block the horizontal transfer of plasmids of the other incompatibility group^71^. A robustness test (see next section) revealed that this default model version behaves similar to another, computationally more demanding model version that considers incompatibility due to interference during plasmid partition as follows: a daughter cell that arises from division of a bacterium that harbored more than one plasmid of the same incompatibility group will not carry more than one plasmid type of each group. To process the loss of incompatible plasmids in the arising daughter cells, it was assumed that each daughter cell has a random combination of plasmids that all belong to different incompatibility groups, but are still originating from the mother cells plasmid load.

#### Segregation

A plasmid type might not be inherited to both daughter cells, even when no other (incompatible) plasmids are present. To process this, each plasmid type that resides in a cell performing cell division has a probability *τ* to be not copied to one of both arising daughter cells (see Table 2). When only a single plasmid type is present, the segregation probability *τ* is equal to the probability that a plasmid-free daughter cell arises. If more plasmid types are present, this probability is much smaller, given by *τ*_*n*_, with *n* as the number of plasmid types that are present in a cell performing division.

### Horizontal plasmid transfer

Plasmids that are conjugative can also be transferred to cells of another cell lineage by horizontal gene transfer. In this submodel, a focal cell attempts to transfer a plasmid copy to a randomly chosen neighboring cell. The overall probability that the focal cell performs a plasmid transfer attempt is given by:

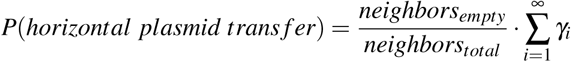

whereas *γ*_*i*_ represents the transfer probability of the *i*-th plasmid that resides on the focal cell; *neighbors*_*total*_ = 8 refers to the local neighbourhood, and *neighbors*_*empty*_ to the number of neighbouring cells which are not occupied.

When the availability of resources is reduced, bacteria can take up and invest less, both in their growth and in the expression of genes such as those required for conjugation. This means that both conjugation and growth are controlled by a local logistic function and not by a global one, which is usually implemented in population models using, for example, ordinary differential equations. This procedure couples plasmid transfer rates to the local growth conditions, which has been reported by various experimental and simulation studies^12,72–74^. Nevertheless, our model neglects that transiently derepression of newly transferred plasmids may initiate a cascade of conjugative transfer events, which has been described as the epidemic spread phenomenon^75,76^.

Considering that *i >* 1, which means that the focal cell bears more than one plasmid type, the single transfer probabilities *γ*_*i*_ are considered to randomly select the *i*-th plasmid for a transfer attempt. The transfer attempt will only be successful, if the receiving, randomly selected, neighboring lattice cell is neither empty nor occupied by a bacterium that already harbors the same plasmid type. In the default model version transfer is also prohibited when the receiving bacterium already bears another plasmid type from the same incompatibility group.

Considering the exemplary conditions for the focal lattice cell depicted in Figure 1-figure supplement 3, *neighbors*_*empty*_ = 3, *γ*_*i*=1_ = 0.21, *γ*_*i*=2_ = 0, resulting in *P*(*plasmid trans f er*) = (3*/*8)(0.71 + 0) ≈ 0.27, which refers in this case to the probability of plasmid *i* = 1 to perform a transfer attempt. The probability that this is successful is further reduced according to the proportion of potential recipients (non-empty, not already bearing the same plasmid type) in the local neighborhood, here denoted as *neighbors*_*pot*.*recipients*_ = 2, which results in *P*(*horizontal plasmid trans f er*_*real*_) = *P*(*horizontal plasmid trans f er*) · (*neighbors*_*pot*.*recipients*_*/neighbors*_*total*_) = 0.27 ∗ (2*/*8) ≈ 0.07.

## Data and Software Availability

Our model, the PlasmidomeIBM, was implemented in the software environment NetLogo^69^. The GitHub repository listed below provides its full code, embedded in a ready-to-run application. This allows it to reproduce our results by running predefined simulation experiments as well as to run simulations with self-defined parameters. Figures were generated with the software environment R^77^ and associated package ‘ggplot2’^78^.

- ‘PlasmidomeIBM’ simulation model source code and app (NetLogo):GitHub (https://github.com/mzwanzig/PlasmidomeIBM/releases/tag/v1.0)

## Acknowledgements

We acknowledge the support of the Open Access Publication Funds of the TU Dresden and thank the Center for Information Services and High Performance Computing (ZIH) at TU Dresden for generous allocations of computer time.

## Figure supplements

**Figure 1-figure supplement 1.**
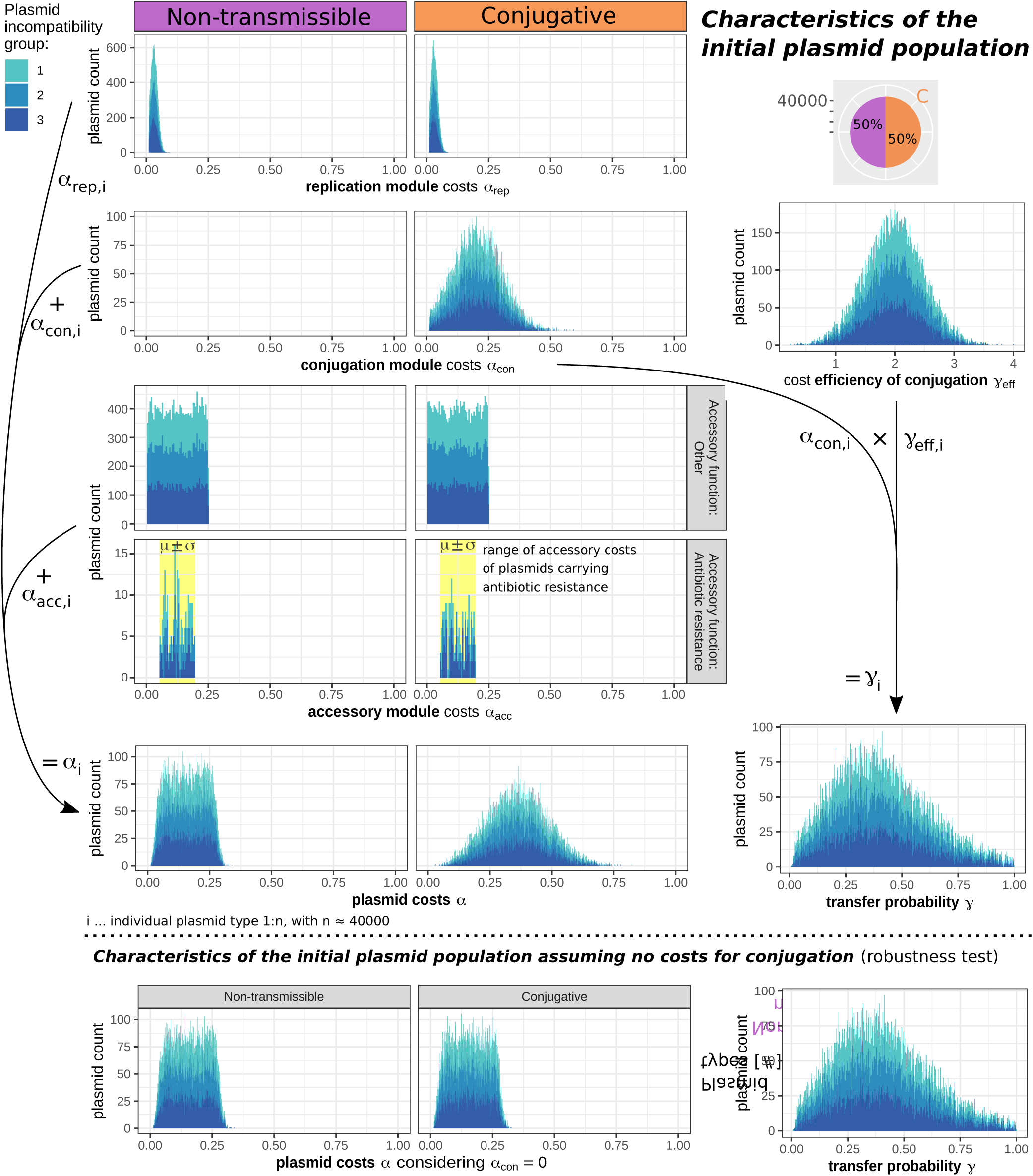
Frequency distribution of plasmid traits in the initial population. Each plasmid is either non-transmissible or conjugative, has some costs *α* = *α*_*rep*_ + *α*_*con*_ + *α*_*acc*_, a transfer probability *γ* = *α*_*con*_ *× γ*_*e f f*_, and, in this example, belongs to one of three incompatibility groups. Some plasmids have nearly no costs for accessory genes, but those that provide antibiotic resistance are assumed to have some average accessory costs *α*_*acc*_. The lower line shows the distribution of plasmid properties used for a robustness test to investigate whether the main results of the model depend on the assumption that conjugation is costly. For this, *α*_*con*_ was set to zero for the calculation of plasmid costs *α*, but not for the calculation of transfer probabilities *γ*. Therefore, non-transmissible plasmids and conjugative plasmids have the same distribution of plasmid costs, while the distribution of transfer probabilities remained the same as in the standard setting above.

**Figure 1-figure supplement 2.**
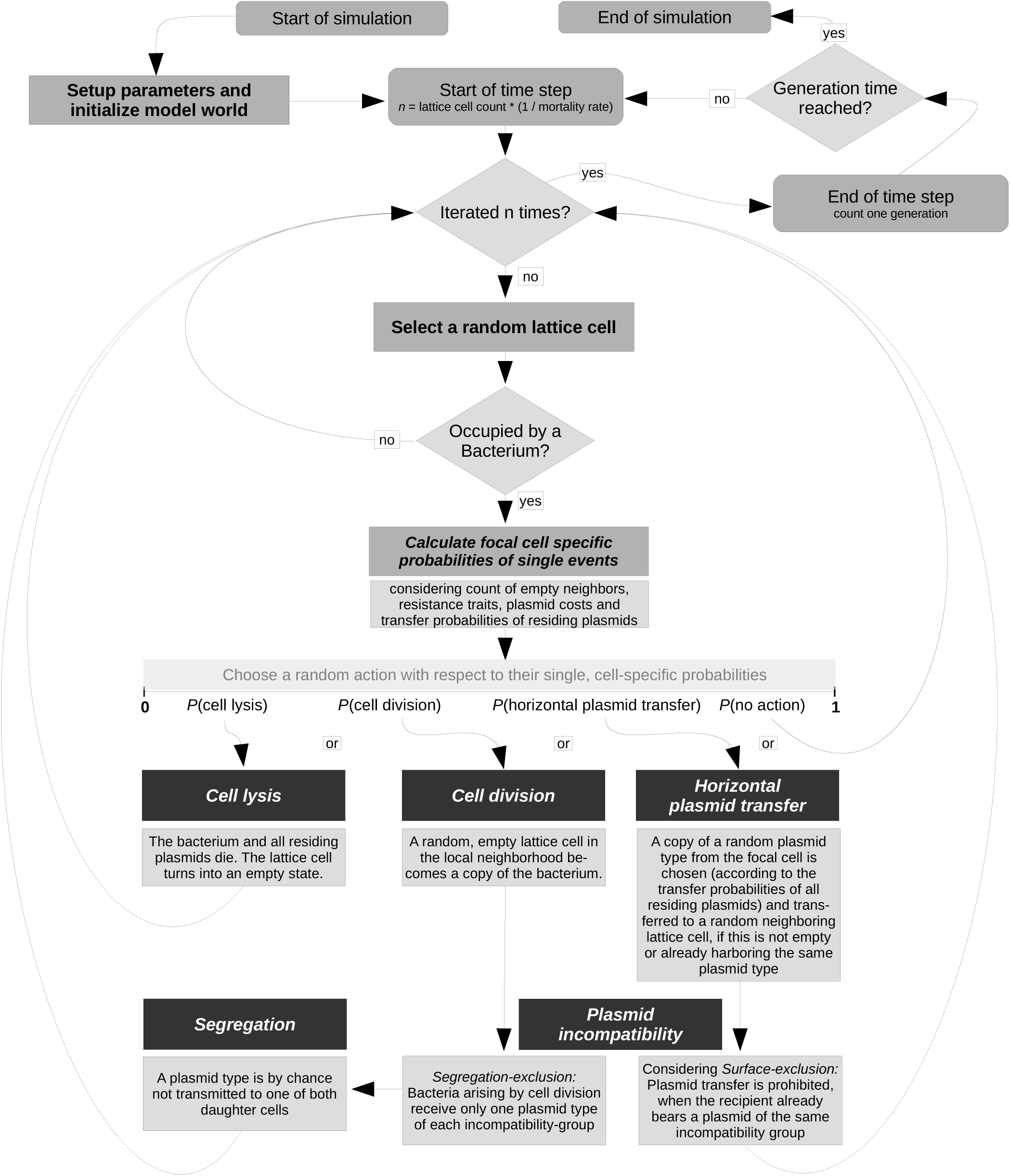
Flow chart of the individual-based plasmidome model. Stochasticity is a major principle in our model: the individual cell that becomes updated is selected at random as well as the action that is chosen regarding to its cell-specific probabilities. Although plasmid types are considered as individuals that can solely be inherited or transfered to other lattice cells, they only perform actions in the context of their host cell environment (the occupied lattice cell (bacterium) they reside on possibly in conjunction with other plasmid types).

**Figure 1-figure supplement fig 3.**
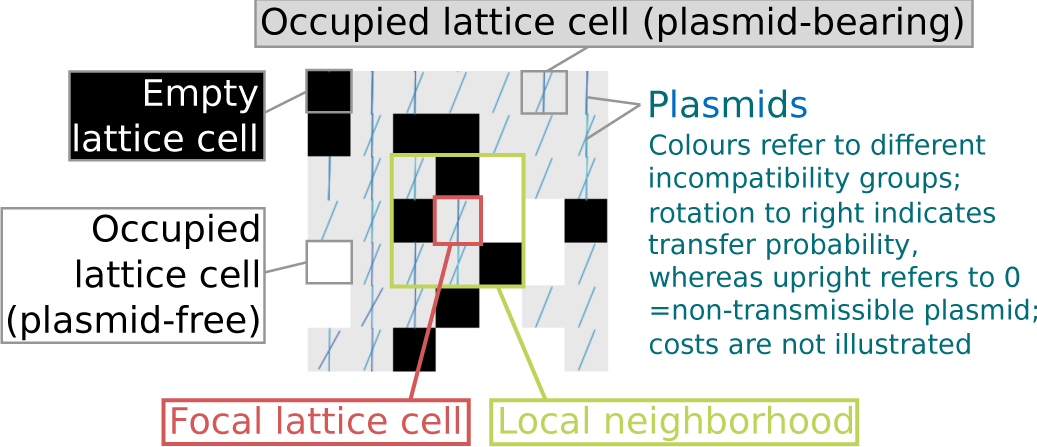
Spatial representation of model entities. The graphic shows a random clipping of the model world, generated by a snapshot from the user interface of NetLogo (the software providing the platform for the development and simulation of the presented individual-based model).

**Figure 2-figure supplement 1.**
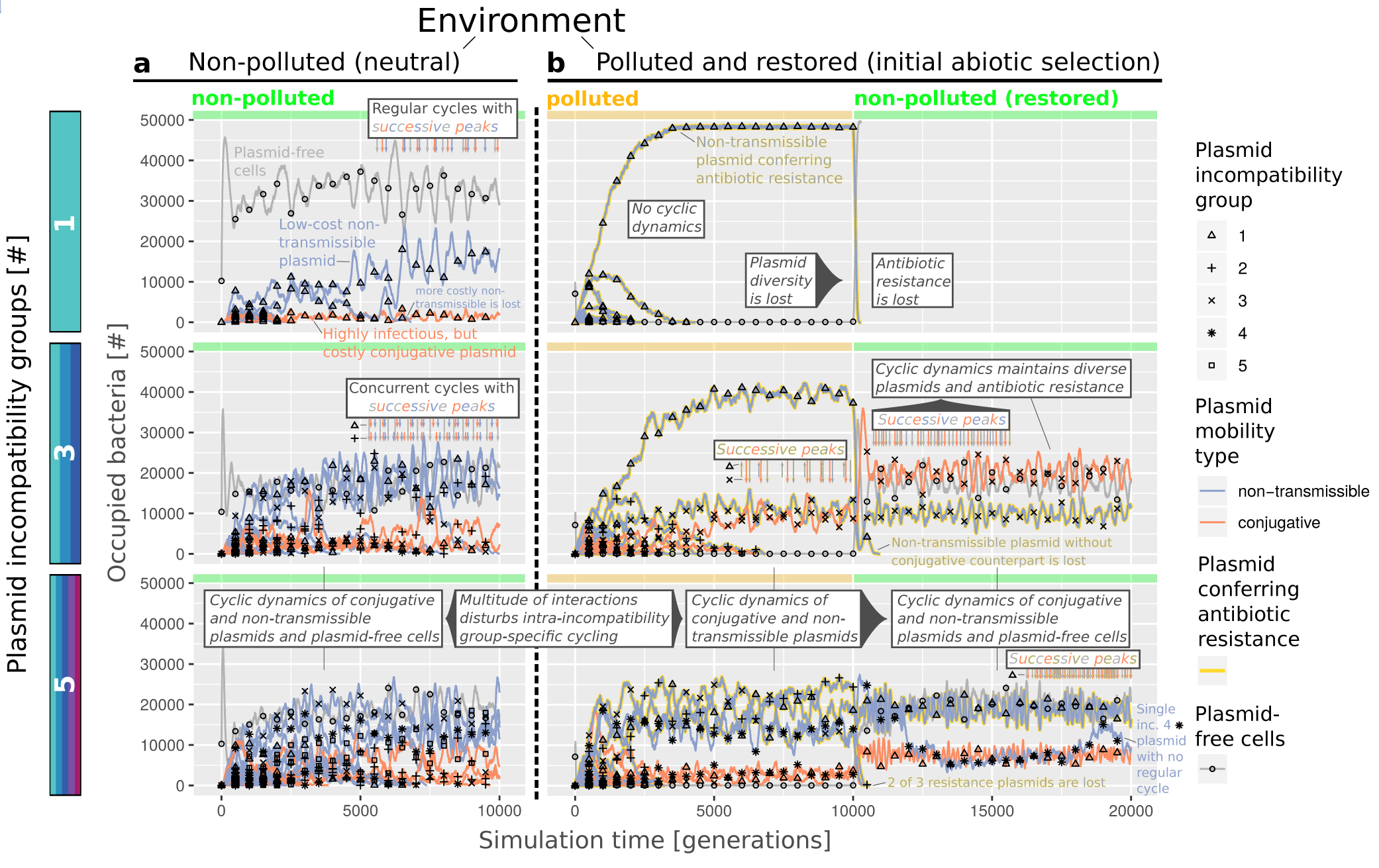
Temporal dynamics of individual plasmid types in non-polluted as well as polluted and restored environments. Each row depicts the results of a single simulation, initialized with a plasmid community consisting either of 1, 3 or 5 different plasmid incompatibility groups considering the absence (‘non-polluted’ environment) or initial presence of antibiotics (for 10000 generations; ‘polluted and restored’ environment). Increases in the frequency of plasmid-free cells are followed by increases in the frequency of conjugative plasmids, to which an increase in the frequency of non-transmissible plasmids of the same incompatibility group occurs. The more incompatibility groups there are, the more this dynamic can be disturbed due to interactions between plasmids of different incompatibility groups and their respective influence on plasmid-free cells.

**Figure 3-figure supplement 1.**
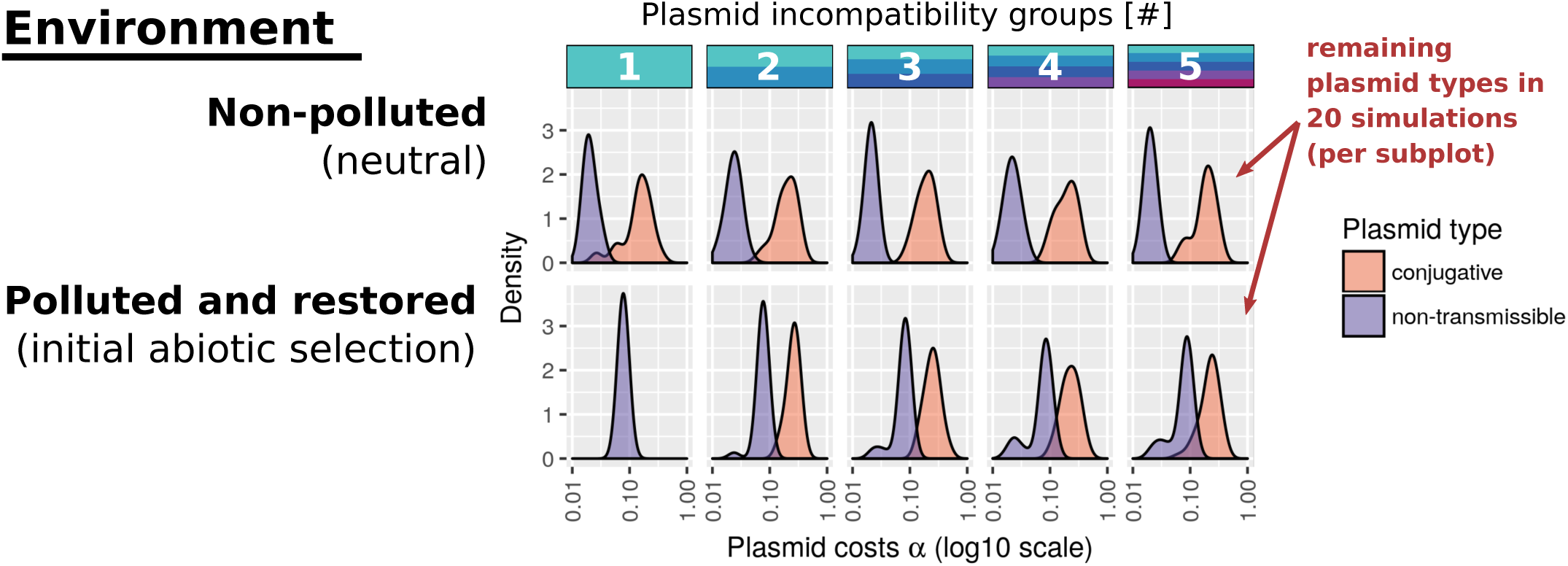
Association between plasmid costs, mobility and the number of incompatibility groups in non-polluted as well as polluted and restored environments. Results refer to the communities that evolved in our simulation experiments considering the absence (‘non-polluted’ environment) or initial presence (for 10000 generations; ‘polluted and restored’ environment) of antibiotics.

**Figure 10-figure supplement 1.**
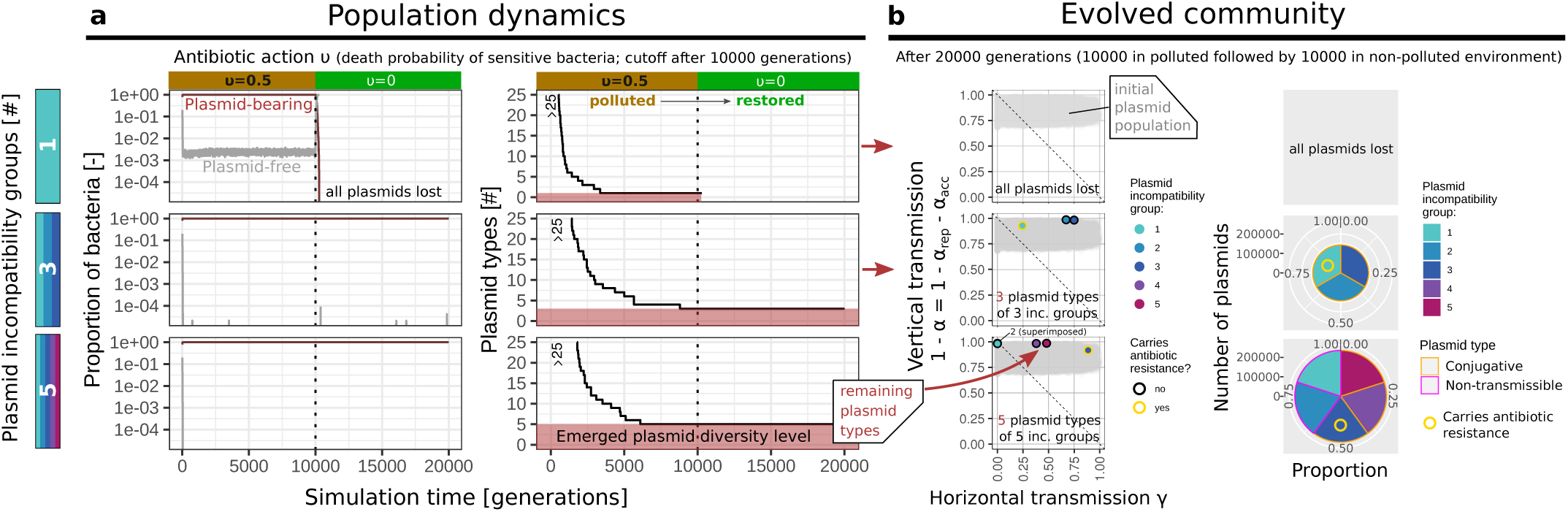
If no costs for conjugation are considered, only one type of plasmid per incompatibility group can survive in a polluted environment that gets restored (initial abiotic selection). Each row depicts the results of a single simulation, initialized with plasmid community consisting either of 1, 3 or 5 different plasmid incompatibility groups.

**Figure 10-figure supplement 2.**
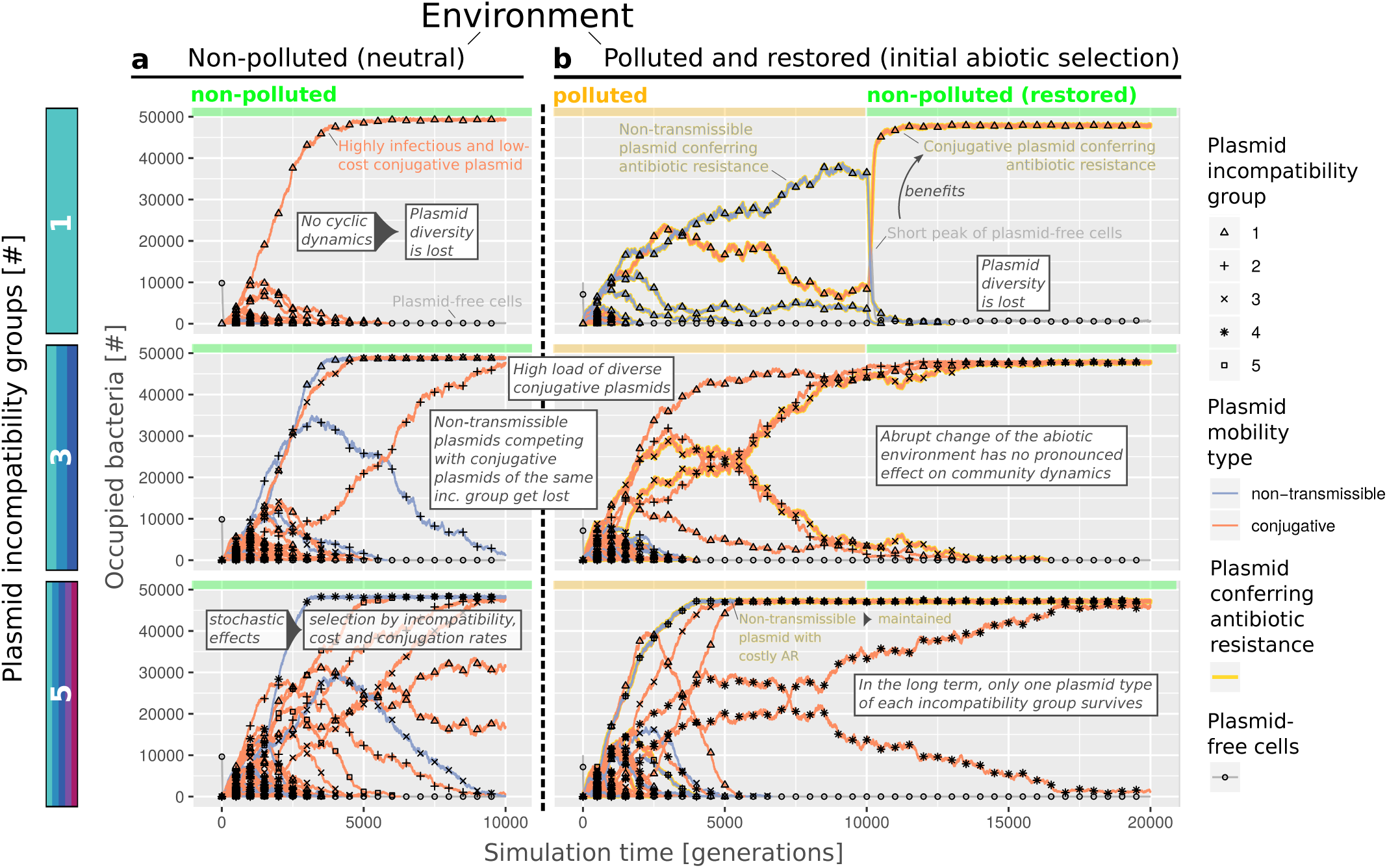
If conjugation is considered to be free of cost, plasmid types of the same incompatibility group are not undergoing intransitive dynamics. Each row depicts the results of a single simulation, initialized with a plasmid community consisting either of 1, 3 or 5 different plasmid incompatibility groups considering the absence (‘non-polluted’ environment) or initial presence (for 10000 generations; ‘polluted and restored’ environment) of antibiotics. In each setting, plasmid-free cells are readily outperformed by highly infectious conjugative plasmids, and only one plasmid type per incompatibility group can survive. Non-transmissible plasmids can survive if all incompatible conjugative plasmids are lost.

## References

1. Touchon, M. et al. Organised genome dynamics in the escherichia coli species results in highly diverse adaptive paths. PLoS Genet. 5, e1000344, DOI: 10.1371/journal.pgen.1000344 (2009).

2. Brockhurst, M. A. et al. The ecology and evolution of pangenomes. Curr. Biol. 29, R1094 – R1103, DOI: https://doi.org/10.1016/j.cub.2019.08.012 (2019).

3. McInerney, J. O., McNally, A. & O’Connell, M. J. Why prokaryotes have pangenomes. Nat. Microbiol. 2, 17040 (2017).

4. Smalla, K. & Sobecky, P. A. The prevalence and diversity of mobile genetic elements in bacterial communities of different environmental habitats: insights gained from different methodological approaches. FEMS Microbiol. Ecol. 42, 165–175, DOI: 10.1111/j.1574-6941.2002.tb01006.x (2002).

5. Boronin, A. M. Diversity of pseudomonas plasmids: To what extent? FEMS Microbiol. Lett. 100, 461–467, DOI: 10.1111/j.1574-6968.1992.tb14077.x (1992).

6. Ricci, G., Borgo, F. & Fortina, M. G. Plasmids from lactobacillus helveticus: distribution and diversity among natural isolates. Lett. Appl. Microbiol. 42, 254–258, DOI: 10.1111/j.1472-765x.2005.01847.x (2006).

7. Brown, C. J. et al. Diverse broad-host-range plasmids from freshwater carry few accessory genes. Appl. Environ. Microbiol. 79, 7684–7695, DOI: 10.1128/AEM.02252-13 (2013).

8. Kwasiborski, A. et al. Core genome and plasmidome of the quorum-quenching bacterium rhodococcus erythropolis. Genetica 143, 253–261, DOI: 10.1007/s10709-015-9827-4 (2015).

9. Dib, J. R., Wagenknecht, M., Farías, M. E. & Meinhardt, F. Strategies and approaches in plasmidome studies—uncovering plasmid diversity disregarding of linear elements? Front. Microbiol. 6, 463, DOI: 10.3389/fmicb.2015.00463 (2015).

10. Medaney, F., Ellis, R. J. & Raymond, B. Ecological and genetic determinants of plasmid distribution in escherichia coli. Environ. Microbiol. 18, 4230–4239, DOI: 10.1111/1462-2920.13552 (2016).

11. Stewart, F. M. & Levin, B. R. The population biology of bacterial plasmids: A priori conditions for the existence of conjugationally transmitted factors. Genetics 87, 209–228 (1977).

12. Merkey, B. V., Lardon, L. A., Seoane, J. M., Kreft, J.-U. & Smets, B. F. Growth dependence of conjugation explains limited plasmid invasion in biofilms: an individual-based modelling study. Environ. Microbiol. 13, 2435–2452 (2011).

13. del Campo, I. et al. Determination of conjugation rates on solid surfaces. A Special Issue on Chromosom. Dyn. Mem. Prof. Kurt Nord. 67, 174–182, DOI: 10.1016/j.plasmid.2012.01.008 (2012).

14. San Millan, A. et al. Positive selection and compensatory adaptation interact to stabilize non-transmissible plasmids. Nat. communications 5, 5208, DOI: 10.1038/ncomms6208 (2014).

15. Stevenson, C., Hall, J. P. J., Brockhurst, M. A. & Harrison, E. Plasmid stability is enhanced by higher-frequency pulses of positive selection. Proc. Royal Soc. Lond. B: Biol. Sci. 285, DOI: 10.1098/rspb.2017.2497 (2018).

16. Harrison, E., Hall, J. P. J. & Brockhurst, M. A. Migration promotes plasmid stability under spatially heterogeneous positive selection. Proc. Royal Soc. B: Biol. Sci. 285, 20180324, DOI: 10.1098/rspb.2018.0324 (2018).

17. Zwanzig, M. et al. Mobile compensatory mutations promote plasmid survival. mSystems 4, DOI: 10.1128/mSystems.00186-18 (2019).

18. Bergstrom, C. T., Lipsitch, M. & Levin, B. R. Natural selection, infectious transfer and the existence conditions for bacterial plasmids. Genetics 155, 1505–1519 (2000).

19. Gelder, L. d., Ponciano, J. M., Joyce, P. & Top, E. M. Stability of a promiscuous plasmid in different hosts: no guarantee for a long-term relationship. Microbiol. (Reading, England) 153, 452–463, DOI: 10.1099/mic.0.2006/001784-0 (2007).

20. Heuer, H., Fox, R. E. & Top, E. M. Frequent conjugative transfer accelerates adaptation of a broad-host-range plasmid to an unfavorable pseudomonas putida host. FEMS Microbiol. Ecol. 59, 738–748, DOI: 10.1111/j.1574-6941.2006.00223.x (2007).

21. Hall, J. P. J. et al. Environmentally co-occurring mercury resistance plasmids are genetically and phenotypically diverse and confer variable context-dependent fitness effects. Environ. Microbiol. DOI: 10.1111/1462-2920.12901 (2015).

22. Hall, J. P. J., Williams, D., Paterson, S., Harrison, E. & Brockhurst, M. A. Positive selection inhibits gene mobilization and transfer in soil bacterial communities. Nat. Ecol. & Evol. 1, 1348–1353 (2017).

23. Cooper, T. F. & Heinemann, J. A. Postsegregational killing does not increase plasmid stability but acts to mediate the exclusion of competing plasmids. Proc. Natl. Acad. Sci. 97, 12643–12648, DOI: 10.1073/pnas.220077897 (2000).

24. Haft, R. J. F., Mittler, J. E. & Traxler, B. Competition favours reduced cost of plasmids to host bacteria. ISME J 3, 761–769 (2009).

25. Cooper, T. F., Tiago, P. & Heinemann, J. A. Within-host competition selects for plasmid-encoded toxin-antitoxin systems. Proc. Royal Soc. B: Biol. Sci. 277, 3149–3155, DOI: 10.1098/rspb.2010.0831 (2010).

26. Silva, R. F. et al. Pervasive sign epistasis between conjugative plasmids and drug-resistance chromosomal mutations. PLOS Genet. 7, 1–10, DOI: 10.1371/journal.pgen.1002181 (2011).

27. Platt, T. G., Morton, E. R., Barton, I. S., Bever, J. D. & Fuqua, C. Ecological dynamics and complex interactions of agrobacterium megaplasmids. Front. Plant Sci. 5, 635, DOI: 10.3389/fpls.2014.00635 (2014).

28. Morton, E. R., Platt, T. G., Fuqua, C. & Bever, J. D. Non-additive costs and interactions alter the competitive dynamics of co-occurring ecologically distinct plasmids. Proceedings. Biol. sciences / The Royal Soc. 281, 20132173, DOI: 10.1098/rspb.2013.2173 (2014).

29. San Millan, A., Heilbron, K. & MacLean, R. C. Positive epistasis between co-infecting plasmids promotes plasmid survival in bacterial populations. The ISME J. 8, 601–612 (2014).

30. Werisch, M., Berger, U. & Berendonk, T. U. Conjugative plasmids enable the maintenance of low cost non-transmissible plasmids. Plasmid 91, 96–104, DOI: 10.1016/j.plasmid.2017.04.004 (2017).

31. Gama, J. A., Zilhão, R. & Dionisio, F. Conjugation efficiency depends on intra and intercellular interactions between distinct plasmids: Plasmids promote the immigration of other plasmids but repress co-colonizing plasmids. Plasmid 93, 6–16, DOI: doi:10.1016/j.plasmid.2017.08.003 (2017).

32. Gama, J. A., Zilhão, R. & Dionisio, F. Multiple plasmid interference – pledging allegiance to my enemy’s enemy. Plasmid 93, 17 –23, DOI: https://doi.org/10.1016/j.plasmid.2017.08.002 (2017).

33. Lopatkin, A. J. et al. Persistence and reversal of plasmid-mediated antibiotic resistance. Nat. Commun. 8, 1689 (2017).

34. Shintani, M., Sanchez, Z. K. & Kimbara, K. Genomics of microbial plasmids: classification and identification based on replication and transfer systems and host taxonomy. Front. Microbiol. 6, 242, DOI: 10.3389/fmicb.2015.00242 (2015).

35. Zhang, T., Zhang, X.-X. & Ye, L. Plasmid metagenome reveals high levels of antibiotic resistance genes and mobile genetic elements in activated sludge. PLOS ONE 6, 1–7, DOI: 10.1371/journal.pone.0026041 (2011).

36. Brown Kav, A. et al. Insights into the bovine rumen plasmidome. Proc. Natl. Acad. Sci. 109, 5452–5457, DOI: 10.1073/pnas.1116410109 (2012). http://www.pnas.org/content/109/14/5452.full.pdf.

37. Jørgensen, T. S., Xu, Z., Hansen, M. A., Sørensen, S. J. & Hansen, L. H. Hundreds of circular novel plasmids and dna elements identified in a rat cecum metamobilome. PLOS ONE 9, 1–9, DOI: 10.1371/journal.pone.0087924 (2014).

38. Luo, W., Xu, Z., Riber, L., Hansen, L. H. & Sørensen, S. J. Diverse gene functions in a soil mobilome. Soil Biol. Biochem. 101, 175 –183, DOI: https://doi.org/10.1016/j.soilbio.2016.07.018 (2016).

39. Shi, Y., Zhang, H., Tian, Z., Yang, M. & Zhang, Y. Characteristics of arg-carrying plasmidome in the cultivable microbial community from wastewater treatment system under high oxytetracycline concentration. Appl. Microbiol. Biotechnol. 102, 1847–1858, DOI: 10.1007/s00253-018-8738-6 (2018).

40. Li, L., Norman, A., Hansen, L. & Sorensen, S. Metamobilomics – expanding our knowledge on the pool of plasmid encoded traits in natural environments using high-throughput sequencing. Clin. Microbiol. Infect. 18, 8 –11, DOI: https://doi.org/10.1111/j.1469-0691.2012.03862.x (2012).

41. Wein, T., Hülter, N. F., Mizrahi, I. & Dagan, T. Emergence of plasmid stability under non-selective conditions maintains antibiotic resistance. Nat. Commun. 10, 2595 (2019).

42. Hall, J. P. J. & Harrison, E. Bacterial evolution: Resistance is a numbers game. Nat. Microbiol. 1, 16235 EP – (2016).

43. Turner, S. L., Bailey, M. J., Lilley, A. K. & Thomas, C. M. Ecological and molecular maintenance strategies of mobile genetic elements. FEMS Microbiol. Ecol. 42, 177–185, DOI: 10.1111/j.1574-6941.2002.tb01007.x (2002).

44. Kothari, A. et al. Large circular plasmids from groundwater plasmidomes span multiple incompatibility groups and are enriched in multimetal resistance genes. mBio 10, DOI: 10.1128/mBio.02899-18 (2019). https://mbio.asm.org/content/10/1/e02899-18.full.pdf.

45. Godoy, O., Bartomeus, I., Rohr, R. P. & Saavedra, S. Towards the integration of niche and network theories. Trends Ecol. & Evol. 33, 287 –300, DOI: https://doi.org/10.1016/j.tree.2018.01.007 (2018).

46. Cairns, J. et al. Black queen evolution and trophic interactions determine plasmid survival after the disruption of the conjugation network. mSystems 3, DOI: 10.1128/mSystems.00104-18 (2018).

47. Amos, G. C. A. et al. Validated predictive modelling of the environmental resistome. ISME J 9, 1467–1476 (2015).

48. Zhou, J. & Ning, D. Stochastic community assembly: Does it matter in microbial ecology? Microbiol. Mol. Biol. Rev. 81, DOI: 10.1128/MMBR.00002-17 (2017). https://mmbr.asm.org/content/81/4/e00002-17.full.pdf.

49. Ben-Hur, E. & Kadmon, R. An experimental test of the area heterogeneity tradeoff. Proc. Natl. Acad. Sci. 117, 4815–4822, DOI: 10.1073/pnas.1911540117 (2020). https://www.pnas.org/content/117/9/4815.full.pdf.

50. Fernández, N., Maldonado, C. & Gershenson, C. Information Measures of Complexity, Emergence, Self-organization, Homeostasis, and Autopoiesis, 19–51 (Springer Berlin Heidelberg, Berlin, Heidelberg, 2014).

51. Kerr, B., Riley, M. A., Feldman, M. W. & Bohannan, B. J. M. Local dispersal promotes biodiversity in a real-life game of rock-paper-scissors. Nature 418, 171–174 (2002).

52. Tecon, R., Ebrahimi, A., Kleyer, H., Erev Levi, S. & Or, D. Cell-to-cell bacterial interactions promoted by drier conditions on soil surfaces. Proc. Natl. Acad. Sci. 115, 9791–9796, DOI: 10.1073/pnas.1808274115 (2018).

53. Grimm, V. & Berger, U. Robustness analysis: Deconstructing computational models for ecological theory and applications. Next generation ecological modelling, concepts, theory: structural realism, emergence, predictions 326, 162–167, DOI: 10.1016/j.ecolmodel.2015.07.018 (2016).

54. Koraimann, G. & Wagner, M. A. Social behavior and decision making in bacterial conjugation. Front. cellular infection microbiology 4, 54, DOI: 10.3389/fcimb.2014.00054 (2014).

55. Smillie, C., Garcillán-Barcia, M. P., Francia, M. V., Rocha, E. P. C. & La Cruz, F. d. Mobility of Plasmids. Microbiol. Mol. Biol. Rev. 74, 434–452 (2010).

56. Heuer, H., Abdo, Z. & Smalla, K. Patchy distribution of flexible genetic elements in bacterial populations mediates robustness to environmental uncertainty. FEMS microbiology ecology 65, 361–371, DOI: 10.1111/j.1574-6941.2008.00539.x (2008).

57. Erkus, O. et al. Multifactorial diversity sustains microbial community stability. The Isme J. 7, 2126–2136 (2013).

58. Soliveres, S. et al. Intransitive competition is common across five major taxonomic groups and is driven by productivity, competitive rank and functional traits. J. Ecol. 106, 852–864, DOI: 10.1111/1365-2745.12959 (2018).

59. Kelsic, E. D., Zhao, J., Vetsigian, K. & Kishony, R. Counteraction of antibiotic production and degradation stabilizes microbial communities. Nature 521, 516 (2015).

60. Hall, J. P. J., Harrison, E. & Brockhurst, M. A. Competitive species interactions constrain abiotic adaptation in a bacterial soil community. Evol. Lett. 2, 580–589, DOI: 10.1002/evl3.83 (2018).

61. Jørgensen, T. S., Kiil, A. S., Hansen, M. A., Sørensen, S. J. & Hansen, L. H. Current strategies for mobilome research. Front. Microbiol. 5, 750, DOI: 10.3389/fmicb.2014.00750 (2015).

62. Stalder, T., Press, M. O., Sullivan, S., Liachko, I. & Top, E. M. Linking the resistome and plasmidome to the microbiome. The ISME J. (2019).

63. Aminov, R. I. Horizontal gene exchange in environmental microbiota. Front. Microbiol. 2, DOI: 10.3389/fmicb.2011.00158 (2011).

64. Klümper, U. et al. Broad host range plasmids can invade an unexpectedly diverse fraction of a soil bacterial community. The ISME J. 9, 934–945 (2015).

65. Garcillán-Barcia, M. P., Alvarado, A. & La Cruz, F. d. Identification of bacterial plasmids based on mobility and plasmid population biology. FEMS Microbiol. Rev. 35, 936–956 (2011).

66. Dahlberg, C. & Chao, L. Amelioration of the cost of conjugative plasmid carriage in eschericha coli k12. Genetics 165, 1641–1649 (2003).

67. Hellweger, F. L., Clegg, R. J., Clark, J. R., Plugge, C. M. & Kreft, J.-U. Advancing microbial sciences by individual-based modelling. Nat Rev Micro 14, 461–471 (2016).

68. Grimm, V. et al. The odd protocol: A review and first update. Ecol. Model. 221, 2760–2768, DOI: 10.1016/j.ecolmodel.2010.08.019 (2010).

69. Wilensky, U. NetLogo. Center for Connected Learning and Computer-Based Modeling, Northwestern University, Evanston, IL. (1999).

70. Hülter, N. et al. An evolutionary perspective on plasmid lifestyle modes. Curr. Opin. Microbiol. 38, 74 –80, DOI: https://doi.org/10.1016/j.mib.2017.05.001 (2017). Mobile genetic elements and HGT in prokaryotes * Microbiota.

71. Humbert, M., Huguet, K. T., Coulombe, F. & Burrus, V. Entry exclusion of conjugative plasmids of the inca, incc and related untyped incompatibility groups. J. Bacteriol. DOI: 10.1128/JB.00731-18 (2019). https://jb.asm.org/content/early/2019/03/06/JB.00731-18.full.pdf.

72. Fox, R. E., Zhong, X., Krone, S. M. & Top, E. M. Spatial structure and nutrients promote invasion of IncP-1 plasmids in bacterial populations. ISME J 2, 1024–1039 (2008).

73. Haagensen, J. A. J., Hansen, S. K., Johansen, T. & Molin, S. In situ detection of horizontal transfer of mobile genetic elements. FEMS Microbiol. Ecol. 42, 261–268, DOI: 10.1111/j.1574-6941.2002.tb01016.x (2002).

74. Seoane, J. et al. An individual-based approach to explain plasmid invasion in bacterial populations. FEMS Microbiol. Ecol. 75, 17–27 (2011).

75. Lundquist, P. D. & Levin, B. R. Transitory derepression and the maintenance of conjugative plasmids. Genetics 113, 483–497 (1986).

76. Ghigo, J. M. Natural conjugative plasmids induce bacterial biofilm development. Nature 412, 442–445, DOI: 10.1038/35086581 (2001).

77. RCoreTeam. R: A Language and Environment for Statistical Computing. R Foundation for Statistical Computing, Vienna, Austria (2019).

78. Wickham, H. ggplot2: elegant graphics for data analysis (Springer-Verlag New York, 2016).

